# An activity-resistance tradeoff constrains enzyme evolution

**DOI:** 10.64898/2026.01.19.700455

**Authors:** Alexander W. Sarkis, Jens Laurids Sørensen, Teis Esben Sondergaard, Kåre Lehmann Nielsen, Jens Christian Frisvad, Douglas L. Theobald, Lizbeth Hedstrom

**Affiliations:** Graduate Program in Biochemistry and Biophysics, Brandeis University, Waltham, Massachusetts, 02454, USA; Department of Chemistry and Bioscience, Aalborg University, Esbjerg 6700, Denmark; Department of Chemistry and Bioscience, Aalborg University, Aalborg 9220, Denmark; Department of Biotechnology and Biomedicine (DTU Bioengineering), Technical University of Denmark, Søltofts Plads, 2800 Kongens Lyngby, Denmark; Department of Biochemistry and Biophysics, Brandeis University, Waltham, Massachusetts, 02454, USA; Department of Biology, Brandeis University, Waltham, Massachusetts, 02454, USA; Department of Chemistry, Brandeis University, Waltham, Massachusetts, 02454, USA

**Keywords:** IMP dehydrogenase, drug resistance, mycophenolic acid, enzyme evolution, biosynthetic gene cluster

## Abstract

The presence of self-resistance genes in antibiotic-producing organisms poses a paradox: how can resistance evolve before the antibiotic exists, and how can an antibiotic producer arise without first evolving resistance? Here we examine the evolutionary origins of self-resistance to mycophenolic acid (MPA), an inhibitor of inosine monophosphate dehydrogenase (IMPDH). The MPA biosynthetic gene cluster (BGC) includes a resistant IMPDH-B. Homologs of IMPDH-B occur not only in MPA producers but also in many non-producing fungi, where remnants of the MPA BGC remain detectable. The phylogeny of IMPDH-B is incongruent with the fungal species tree, consistent with multiple horizontal gene transfer events between Aspergillus and Sordariomycetes. We characterized eleven extant IMPDH-Bs, five from MPA producers and six from nonproducers, along with seven resurrected ancestral enzymes (Anc1–Anc7). MPA resistance appeared between Anc2 and Anc3 and coincided with a loss of catalytic efficiency. Across both ancestral and extant enzymes, MPA resistance correlated strongly with reduced activity, revealing a robust activity–resistance trade-off that has persisted for millions of years. Unexpectedly, both the IMPDH-Bs and ancestral enzymes Anc3–Anc7 were also resistant to ribavirin-5′-monophosphate (RVP), an IMP-competitive inhibitor. Because MPA and RVP bind to similar enzyme conformations, the activity-resistance trade-off likely reflects a design constraint imposed by the need to maintain resistance to multiple inhibitors. Intriguingly, although Anc1 and Anc2 are equally sensitive to MPA, Anc2 shows reduced susceptibility to RVP. This pattern suggests that pre-existing resistance to another IMPDH inhibitor may have created a permissive background for the later evolution of MPA biosynthesis.

**Significance:** Antibiotic producers must be resistant to the toxins that they produce, but how such self-resistance develops is a mystery. The mycophenolic acid (MPA) biosynthetic gene cluster (BGC) encodes a resistant variant of the MPA target IMPDH (IMPDH-B). Many fungi retain IMPDH-B although they have lost the ability to produce MPA. The IMPDH-B and species phylogenies are incongruent, suggesting evolution of the BGC was complicated. MPA resistance correlates with low catalytic efficiency in modern and ancestral IMPDHs, revealing a robust design constraint tradeoff. Surprisingly, IMPDH-Bs are also resistant to an IMP-competitive inhibitor (RVP). RVP resistance appears to have emerged before MPA resistance. Perhaps resistance to RVP created a background that permitted the genesis of a new toxin.

---

Fungi employ a wide variety of self-resistance mechanisms to enable the biosynthesis of toxic natural products (1–4). How these self-resistance mechanisms evolved is a classic chicken-and-egg problem: a toxic molecule cannot be produced in the absence of self-resistance, yet how does self-resistance develop in the absence of the toxic molecule? The biosynthesis of the fungal natural product mycophenolic acid (MPA) exemplifies this issue (Figure 1A). MPA and its prodrug mycophenolate mofetil are immunosuppressive agents used in organ transplants and to treat autoimmune diseases (5, 6). MPA is also a common environmental antimicrobial and a frequent contaminant of food and silage (7–9). *Penicillium brevicompactum* is the industrial MPA producer, though several other *Penicillium* and Aspergillus species and a few other filamentous fungi also produce MPA (7, 10–24). The target of MPA is inosine monophosphate dehydrogenase (IMPDH), the rate-limiting enzyme in guanine nucleotide biosynthesis (Figure 1B) (25). IMPDH is also the target of the immunosuppressive mizoribine (a natural product also produced by *Penicillium*), oxanosine (a natural product produced by *Streptomyces capreolus*) as well as the antiviral ribavirin (Figure 1C) (26, 27). These nucleosides are converted to monophosphates that bind in the IMP site, while MPA binds in the NAD^+^ cofactor site. The MPA biosynthetic gene cluster (BGC) includes the self-resistance gene *mpaF* that encodes an MPA-resistant (MPA^R^) IMPDH (IMPDH-B; IMPDH-A denotes the “normal” MPA-sensitive (MPA^S^) enzyme) (28, 29). *P. brevicompactum* IMPDH-B (*Pbre*B) is approximately 500-fold more resistant to MPA than a typical eukaryotic IMPDH even though the active sites and the MPA binding sites are conserved in MPA^R^ and MPA^S^ enzymes (Figure 1D) (30, 31). Curiously, most *Penicillium* and several other fungi in the Eurotiales order contain both IMPDH-A and IMPDH-B, regardless of whether they produce MPA.

**Figure 1.**
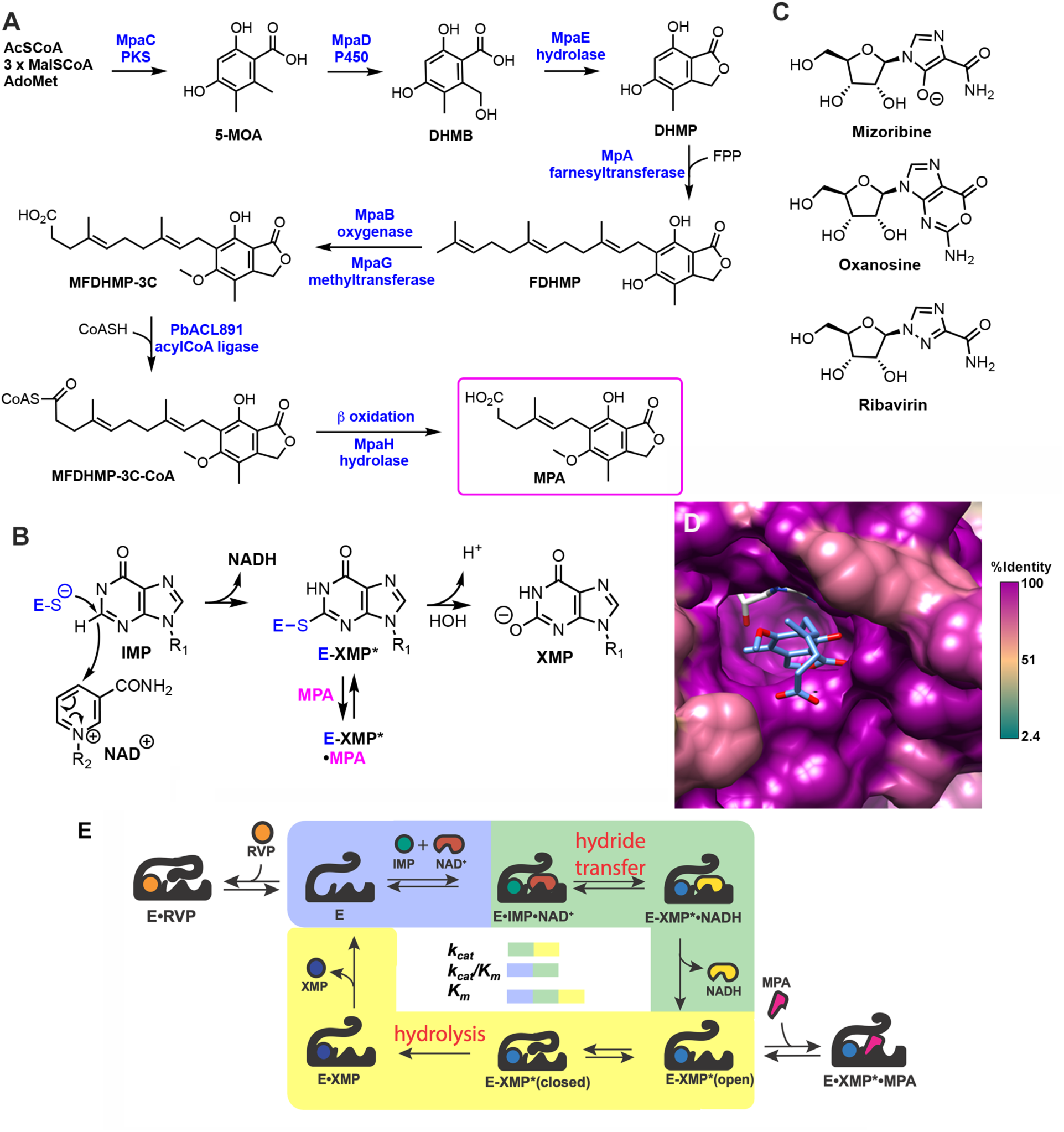
MPA inhibits IMPDH. **A.** Biosynthesis of MPA. MpaC is a polyketide synthase (PKS) that catalyzes the synthesis of the phthalide group. Subsequent MpaDE-catalyzed hydroxylation and lactonization produces 3,5-dihydroxy-6-methylphthalide (DHMP), which is converted to 4-farnesyl-3,5-dihydroxy-6-methylphthalide (FDHMP) by the action of the farnesyl transferase MpaA. The methoxy group is installed by the oxygenase MpaB and O-methyltransferase MpaG to produce 5-O-methyl-FDHMP (MFDHMP). MFDHMP is converted to the CoA ester by PbACL891, an enzyme not encoded in the MPA BGC. Finally, the farnesyl tail is processed by β-oxidation and the CoA group is removed by acyl-CoA hydrolase MpaH to produce MPA. **B.** The chemical transformations of the IMPDH reaction. R_1_ = ribose-5’-monophosphate; R_2_ = adenosyl-5’-diphosphate. **C.** Structure of IMPDH inhibitors. **D.** The structure of the E-XMP*•MPA complex of *Cricetulus griseus* IMPDH2 (PDB: 1JR1) (78) is colored by conservation in an alignment of 83 fungal IMPDHs, including 62 IMPDH-As (MPA-sensitive) and 21 IMPDH-Bs (MPA-resistant) plus the human and *C. griseu*s IMPDH2s (MPA-sensitive). XMP* is white, MPA is cornflower blue. Residues that interact with MPA in *C. griseu*s IMPDH2 are conserved, including D274, S275, S276, N303, C331, R332, T333, and Q441 (*C. griseu*s numbering) (78). This figure was rendered in UCSF Chimera (79). **E.** The kinetic mechanism of IMPDH (25). The value of *k_cat_* includes all the steps after the ternary complex E•IMP•NAD^+^ (blue and green shading) while the value of *k_cat_/K_m_* includes all the steps up to and including the first irreversible step: substrate binding, hydride transfer and NADH release (the first irreversible step under initial rate conditions; blue and green shading). In contrast, the value of *K_m_* includes all the steps in the catalytic cycle (blue, green and yellow).

The remarkable resistance of *Pbre*B derives from changes in kinetics rather than drug affinity (31). MPA traps the E-XMP* covalent intermediate after NADH departs, preventing a conformational change required for the hydrolysis reaction that produces XMP (Figure 1E). The affinity of MPA thus depends on the accumulation of E-XMP* in addition to interactions at the cofactor site. The formation of E-XMP* is rate-limiting in “normal” IMPDHs, so E-XMP* is the predominant enzyme form during catalysis. In contrast, hydrolysis is not rate-limiting in *Pbre*B, so E-XMP* fails to accumulate, making the enzyme resistant to MPA (31).

The structural basis of *Pbre*B resistance has not been fully elucidated (30). *Pbre*B does not resemble prokaryotic IMPDHs that are similarly resistant to MPA (32) and does not contain substitutions that are associated with MPA-resistance in other eukaryotic IMPDHs, e.g., A249T, A249S, Q277R or A462T (human IMPDH2 numbering) (33–36). IMPDH-As and Bs are ∼85% identical, with ∼30 substitutions in the catalytic domain. *Pbre*A becomes 7-fold more resistant to MPA when its C-terminal 30 residues are swapped for those of *Pbre*B. This segment includes half of the binding site for the activating monovalent cation (30). Additional structural determinants of resistance remain to be identified.

Our initial investigations of the origin of IMPDH-Bs suggested that they evolved from a gene duplication that occurred at or before the *Penicillium* genus partitioned into subgenera *Aspergilloides* and *Penicillium* approximately 73.6 MYA (28, 30, 37, 38). IMPDH gene amplification provides MPA resistance in other organisms (39–42), so this duplication event could have created a permissive environment for the emergence of MPA biosynthesis, or possibly even been in response to the presence of MPA. Our initial IMPDH-B phylogenetic tree is very similar to those of IMPDH-A and tubulin, suggesting that all three genes evolved under similar fitness pressure (30). Curiously, MPA producers and nonproducers are interspersed throughout these cladograms, suggesting multiple instances of BGC loss or horizontal gene transfer. Unfortunately, at that time, only one genome sequence was available from a MPA nonproducer *Penicillium (P. chrysogenum,* now recognized as *P. rubens Wisconsin* (43)), so the possibility that nonproducers harbored silent MPA BGCs could not be eliminated in most cases. *P. rubens Wisconsin* is more resistant to MPA than the non-IMPDH-B containing *A. nidulans*, suggesting that its IMPDH-B (*PruW*B) might be MPA-resistant (30). Characterization of recombinant *PruW*B failed to support this hypothesis, although this enzyme was unstable and could only be partially purified (30, 31). Therefore the relationship of IMPDH-Bs to MPA resistance in nonproducers is uncertain.

The increased availability of whole genome sequences from filamentous fungi motivated our reinvestigation of the evolution of IMPDH-Bs and MPA resistance. The resulting IMPDH-B cladogram is not congruent with the fungal species tree. We resurrected 7 ancestors and characterized 11 extant IMPDH-Bs, including enzymes from both MPA producers and nonproducers. All extant IMPDH-Bs are MPA^R^ but less active than MPA^S^ enzymes, regardless of whether they are from MPA producers or nonproducers. Moreover, MPA affinity correlated with catalytic efficiency, revealing a robust design tradeoff. Surprisingly, ancestral and modern IMPDH-Bs are also resistant to the IMP analog RVP, suggesting that MPA-resistant IMPDH-Bs could have originally evolved in response to another IMPDH inhibitor.

## RESULTS

### MPA BGC synteny suggests a complicated evolutionary history

We identified *mpaF* in the genomes of 36 filamentous fungi and then searched for the 8 other genes of the MPA BGC, *mpaA*-*E*, *mpaG*-*H*, and *pbACL891*. MPA BGCs were identified in 15 fungi, including the BGCs from *P. brevicompactum*, *P. roqueforti* and *P. arizonense* that were known at the time of our search (29, 37, 44) (Figure S1A). Seven species were not known to be MPA producers, including *P. egyptiacum*, *P. psychrosexuale*, *P. samsonianum* and four species outside subgenus *Pencillium*: *A. brunneus*, *Byssochlamys sp. AF001*, and two Sordariomycetes fungi, *Bertia moriformis* and *Metarhizium humberi.* Papon and colleagues recently confirmed MPA production in *P. samsonianum* (45).

We inspected the synteny of the MPA BGCs to further assess the evolution of MPA biosynthesis and resistance. Unfortunately, the *P. carneum* BGC was found in non-overlapping contigs, so gene order could not be defined. The BGCs of the 6 *Penicillium* subgenus species, *P. bialowiezense*, *P. brevicompactum*, *P. egyptiacum, P. psychrosexuale*, *P. roqueforti*, and *P. samsonianum,* were colinear while the BGC of *P. parvum* (*Penicillium* subgenus *Aspergilloide)* was very similar (Figure S1A), suggesting MPA biosynthesis was inherited from a common *Penicillium* ancestor. Gene order was conserved in the two *Aspergillus* BGCs, but distinct from the *Penicillium* BGCs. An *mpaB* ortholog could not be identified in either *Aspergillus* BGCs. The gene order of the *Paecilomyces niveus* BGC was distinct from both the *Penicillium* and *Aspergillus* BGCs. These observations are consistent with the hypothesis that IMPDH duplication and MPA production may have emerged prior to the radiation of *Penicillium*, as is the near exclusivity of MPA production to Eurotiomycetes (28, 37, 45).

We also found evidence of likely horizontal gene transfer. The *B. moriformis* BGC was nearly identical to those from *Aspergillus*, with 99.84% identity and only 20 gaps relative to the *A. brunneus* BGC, and 99.06% identity and 184 gaps relative to the *A. pseudoglaucus* BGC. These observations strongly suggest *B. moriformis* obtained the MPA BGC via horizontal gene transfer from an *Aspergillus*. These three BGCs lack *mpaB*, although possible orthologs could be identified in other contigs in *A. brunneus* and *B. moriformis* (40% coverage, 67% identity for both).

Several fungi harbor partial MPA BGCs that are likely involved in the biosynthesis of related natural products, as previously reported for *P. arizonense* CBS141311^T^ (44). Notably, *P. arizonense* HEWt1 is an MPA producer (21), demonstrating that remodeling of the BGC is ongoing. The *Byssochlamys sp. AF001* contains a partial MPA BGC comprised of *mpaC, mpaD, mpaE and mpaF* and an *mpaA* fragment (35% coverage, 71% identity match) that likely produces the phthalide head group. With the exception of *mpaA*, these genes contain frameshifts (45). The gene order of this partial cluster is similar to that of the closely related *Pa. niveus*. The *M. humberi* BGC lacks *mpaG* and *mpaH*, suggesting that either orthologous genes are located elsewhere in the genome or perhaps *M. humberi* only produces the farnesyl-DHMP intermediate. Taken together, these observations suggest the evolutionary history of the MPA BGC includes vertical and horizontal transfer with rearrangement and frequent loss.

### MPA Non-Producer Fungi May Contain Remnants of MPA Biosynthesis

No *mpaA* (prenyltransferase) or *mpaC* (polyketide synthase) candidates with >40% coverage could be identified anywhere in the genomes of the remaining 17 fungi with *mpaF*, indicating that these fungi are unlikely to be MPA producers. However, possible remnants of other MPA biosynthetic genes could be identified in 12 of these genomes (Figure S1B). *P. expansum* contained a putative *mpaH* (hydrolase) with 99% coverage and 73% identity on the same contig as *mpaF* while *P. chrysogenum* contained a putative *mpaE* (hydrolase) (99% coverage, 78% identity) on the *mpaF* contig. The remaining genomes contained *mpaG* candidates (>94% coverage), although none were on the same contig as *mpaF*. These observations suggest that modern nonproducers may have descended from MPA producers and are still in the process of losing the MPA biosynthetic genes via genetic drift.

### IMPDH-Bs Have a Different Evolutionary History than IMPDH-As

A maximum likelihood phylogeny was built with 83 total IMPDH sequences, including *Byssochlamys* and *Talaromyces* (8), *Aspergillus* (28), *Penicilliopsis* (1), *Penicillium* (37) and Eurotiomycetes outgroup fungi (9). These sequences included 21 IMPDH-Bs from 19 species: *Byssochlamys* (2), *Aspergillus* (2), and *Penicillium* (15). Unfortunately, the *mpaF* sequences were not available from *A. brunneus, P. psychrosexuale*, *P. egyptiacum* and *P. bialowiezense* at the time this tree was constructed. The phylogeny strongly indicates that IMPDH-As and IMPDH-Bs are in distinct clades (Figure 2A), as has been concluded from smaller trees (30, 45). As expected for a housekeeping enzyme, IMPDH-A phylogeny is congruent with the previously reported fungal species trees (38, 46). In contrast, IMPDH-B phylogeny bears little resemblance to the species tree, as illustrated by the close relationship of *Pbre*B and *Pnal*B compared to *Pbre*A and *Pnal*A. Interestingly, MPA producers are interspersed with non-producers throughout the tree, showing that MPA production has little correlation with IMPDH-B phylogeny. IMPDH-Bs from MPA producers are clustered at the top of the tree (*A. pseudoglaucus*, *Bysspchlamys spp AF001*, *Pa. niveus* and *P. brevicompactum*) and at the bottom (*P. roqueforti* and *P. samsonianum*). These observations indicate that, like the MPA BGC, IMPDH-Bs have a complicated evolutionary history distinct from that of fungal species, suggesting horizontal gene transfer events.

**Figure 2.**
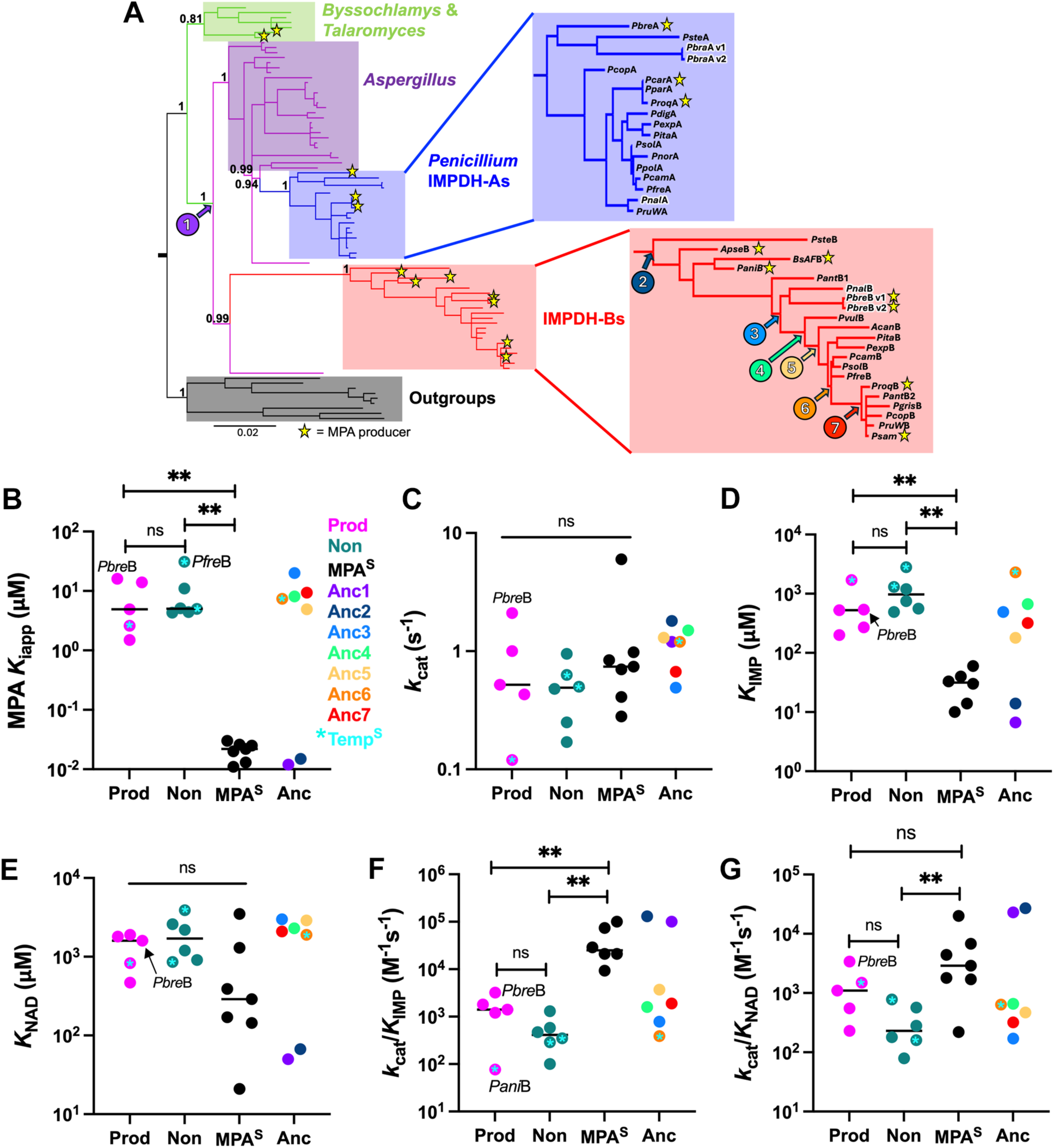
Fungal IMPDH phylogeny, MPA resistance and enzymatic activity. **A.** Phylogenetic tree of 83 Eurotiale order fungal IMPDHs, including 62 IMPDH-As and 21 IMPDH-Bs constructed using BALi-Phy (80). Branches with ≥80% support are shown. Five major subclades are found: *Byssochlamys* and *Talaromyces* (olive green), *Aspergillus* (purple), *Penicillium* IMPDH-As (blue), IMPDH-Bs (red), and outgroups composed closely related *Trichophyton*, *Microsporum*, and *Coccoidiodes.* Bi-partition branch supports are listed to the left of their node of origin. PbreA, PnalA, PbreB and PnalB are highlighted to illustrate the incongruence of the IMPH-A and IMPDH-B phylogeny. Resurrected ancestors are denoted by the number along the branches. MPA producers are denoted with stars. Branch length scale is shown at the bottom. Abbreviations: *Acan*, *Aspergillus candidus*; *Apse*, *Aspergillus pseudoglaucus*; *BsAF*, *Byssochlamys sp* AF001B; *Pani*, *Paecilomyces nivea*; *Pant*, *Penicillium antarcticum*; *Pbre*, *P. brevicompactum*; *Pbra, P. brasilianum*; *Pcam*, *P. camemberti*; *Pcar, P. carneum*; *Pcop, P. coprophilum*; *Pdig, P. digitatum*; *Pexp*, *P. expansum*; *Pfre*, *P. freii*; *Pgris*, *P. griseofulvum*; *Pita*, *P. italicum*; *Pnal*, *P. nalgiovense*; *Pnor, P. nordicum*; *Ppar, P. parvum*; *Ppol, P. polonicum*; *Proq*, *P. roqueforti*; *PruW, P. rubens Wisconsin*; *Psam*, *P. samsonianum*; *Psol*, *P. solitum*; *Pste*, *P. steckii*; *Pvul*, *P. vulpinum*. **B-G.** The values of *K_iapp_* for MPA (**B**), *k_cat_* (**C**), *K_IMP_* (**D**), *K_NAD_* (**E**), *k_cat_/K_IMP_* (**F**) and *k_cat_/K_NAD_* (**G**) are shown for IMPDH-Bs from producers (Prod, magenta) and nonproducers (Non, teal) and ancestors (Anc, colored as noted), as well as MPA-sensitive IMPDHs from the literature (MPA^S^, black, defined as *K_iapp_* ≤ 30 nM), See Figures S3, S4 and S7 and Table S1 and S2 for data and parameters for each enzyme. **, p ≤ 0.01. **, p ≤ 0.01. **B.** IMPDH-Bs and Anc3-Anc6 are MPA-resistant while Anc1 and Anc2 are MPA^S^. **C.** The values of *k_cat_* are similar for IMPDH-Bs and MPA^S^ enzymes. **D.** The values of *K_IMP_* are larger for IMPDH-Bs than MPA^S^ enzymes. **E.** The values of *K_NAD_* are comparable among IMPDH-Bs and MPA^S^ enzymes. **F.** The values of *k_cat_/K_IMP_* of MPA^S^ IMPDHs are greater than IMPDH-Bs. Producer IMPDH-Bs tend to be more active than nonproducer enzymes (p = 0.18; p= 0.02 if the thermolabile *Pani*B value is omitted). **G.** The values of *k_cat_/K_NAD_* of MPA^S^ IMPDHs are greater than nonproducer IMPDH-Bs.

### Modern IMPDH-Bs are MPA-resistant and slow

We characterized 11 extant IMPDH-Bs, 5 from MPA producers and 6 from nonproducers (enzymes are denoted as exemplified by *Pbre*B for *P. brevicompactum* IMPDH-B) (Figures S2-4, Table S1). We also characterized IMPDH-A from *P. brevicompactum* (*Pbre*A); the values of kinetic parameters for *Pbre*A were consistent with previous reports (Table S1). We compared the catalytic activities of these enzymes with MPA^S^ eukaryotic IMPDHs reported in the literature, defined as *K*_iapp_ ≤ 30 nM. All of the IMPDH-Bs were catalytically active (Figure 2B-G, Table S1), including *PruW*B, which was previously reported as inactive (31). *Pbre*B was also more active than previously reported (31). We attribute these discrepancies to an improved purification protocol. Three enzymes (*Pani*B, *Pcam*B and *Pfre*B), all from fungi that live in cold environments, were labile at 25°C, and were assayed at lower temperatures (note that psychrophilic and mesophilic enzymes generally have similar activities at their operational temperatures (47)). All IMPDH-Bs were MPA^R^, regardless of whether they were from producers or non-producers, with values of *K_iapp_* ranging from 1.5 to 31 μM, on average 250-fold higher than MPA^S^ enzymes (Figure 2B). MPA resistance did not vary between producer and non-producer enzymes.

The values of *k_cat_* of IMPDH-Bs and MPA^S^ enzymes were similar (Figure 2C). However, this similarity masks dramatic differences in the underlying kinetic mechanisms (31). Hydrolysis is rate-limiting for MPA^S^ enzymes like hIMPDH2 (Figure 1C,E), so E-XMP* is the predominant enzyme form (48, 49). Since MPA binds to E-XMP*, rate-limiting hydrolysis promotes inhibitor binding. In contrast, hydride transfer is rate-limiting for *Pbre*B, so E-XMP* does not accumulate, making the enzyme MPA^R^ (31).

The values of *K_IMP_* were higher by ≥16-fold for IMPDH-Bs relative to MPA^S^ enzymes (average values of *K_IMP_* are 525, 975 and 32 μM for producer, nonproducer and MPA^S^ enzymes, respectively; Figure 2D). The values of *K*_IMP_ tended to be lower for IMPDH-Bs from producers than nonproducers, although these differences were not significant at p < 0.05 (p = 0.18). The values of *K*_NAD_ also tended to be higher in IMPDH-Bs than MPA^S^ enzymes, but this difference also failed to reach p < 0.05 due to the wide variance of values in MPA^S^ enzymes (Figure 2E). No differences were observed in the values of *K*_NAD_ for IMPDH-Bs from both producers and nonproducers. Unfortunately, these observations also provide little insight into the underlying mechanistic features responsible for MPA-resistance because *K_IMP_* and *K_NAD_* are complex kinetic constants that include all the steps of the catalytic cycle (Figure 1E)(50, 51).

In contrast to *k_cat_* and *K_M_*, the value of *k*_cat_/*K*_M_ includes only the steps up to and including the first irreversible step, i.e., substrate binding, hydride transfer and NADH release in the case of IMPDH. Therefore *k*_cat_/*K*_IMP_ and *k*_cat_/*K*_NAD_ report on the production of E-XMP*, the enzyme form that binds MPA (Figure 1E). IMPDH-Bs were less active than MPA^S^ enzymes as measured by the values of *k*_cat_/*K*_IMP_ (Figure 2F). IMPDH-B-s were also less active MPA^S^ enzymes as measured by the values of *k*_cat_/*K*_NAD_, although this difference did not reach significance with p < 0.05 for producers (p = 0.15)(Figure 2G). *Pbre*B was the most active of all IMPDH-Bs in terms of both *k*_cat_/*K*_IMP_ and *k*_cat_/*K*_NAD_, perhaps reflecting additional optimization required to support the industrial production of MPA. Interestingly, producer IMPDH-Bs tended to be more active than those from nonproducers, although these differences did not reach significance with p < 0.05 (p = 0.18 and 0.08 for *k*_cat_/*K*_IMP_ and *k*_cat_/*K*_NAD_, respectively). The difference between producer and nonproducer *k*_cat_/*K*_IMP_ values is significant if the thermolabile *Pani*B value is excluded (p = 0.019). Therefore values of *k*_cat_/*K*_IMP_ greater than 10^3^ M^−1^s^−1^ may indicate MPA production.

### MPA-resistance emerged between Anc2 and Anc3

IMPDH-B proteins display ≥85% sequence pairwise identity with no sequence gaps (Figure S5). A similar level of conservation is observed between IMPDH-B and -A proteins, although IMPDH-Bs are ∼20 residues shorter. The differences include an ∼15 residue truncation of the N-terminus, a single residue deletion in the junction between the regulatory domain and the catalytic domain and a 5-7 residue deletion in an active site loop. This conservation enabled the resurrection of ancestors with high confidence, with the average posterior probabilities all ≥98% from a tree generated with branch supports of ≥80% (Figure 2A).

We resurrected 7 ancestral enzymes, including the last common ancestor of IMPDH-As and -Bs (Anc1) and the ancestor after the radiation of the IMPDH-B clade (Anc2) (Figure 2A). These two enzymes differ by 24 substitutions (Figure S6). Anc1 and Anc2 are MPA^S^ and display low *K_IMP_* and high *k_cat_/K_IMP_* values characteristic of typical eukaryotic IMPDHs (Figures 2-G and 3A). Therefore the distinct functional characteristics of IMPDH-Bs, and presumably MPA biosynthesis, emerged after Anc2.

We were especially interested in ancestors that spawned both MPA producers and nonproducers (Anc3-7, Figure 2A). Anc3 displays all the characteristics of an IMPDH-B: high *K_iapp_* for MPA (20 μM) accompanied low values of both *k_cat_/K_IMP_* and *k_cat_/K_NAD_* (780 and 170 M^−1^s^−1^, respectively; Figures 2B-G and 3A). Anc2 and Anc3 are 85% identical, differing at 77 residues (Figure S6). These substitutions include the changes in the C-terminal segment already associated with MPA resistance (30). None of these substitutions are in the MPA binding site, but many are in residues that interact with the MPA binding site (Figure 3B).

**Figure 3.**
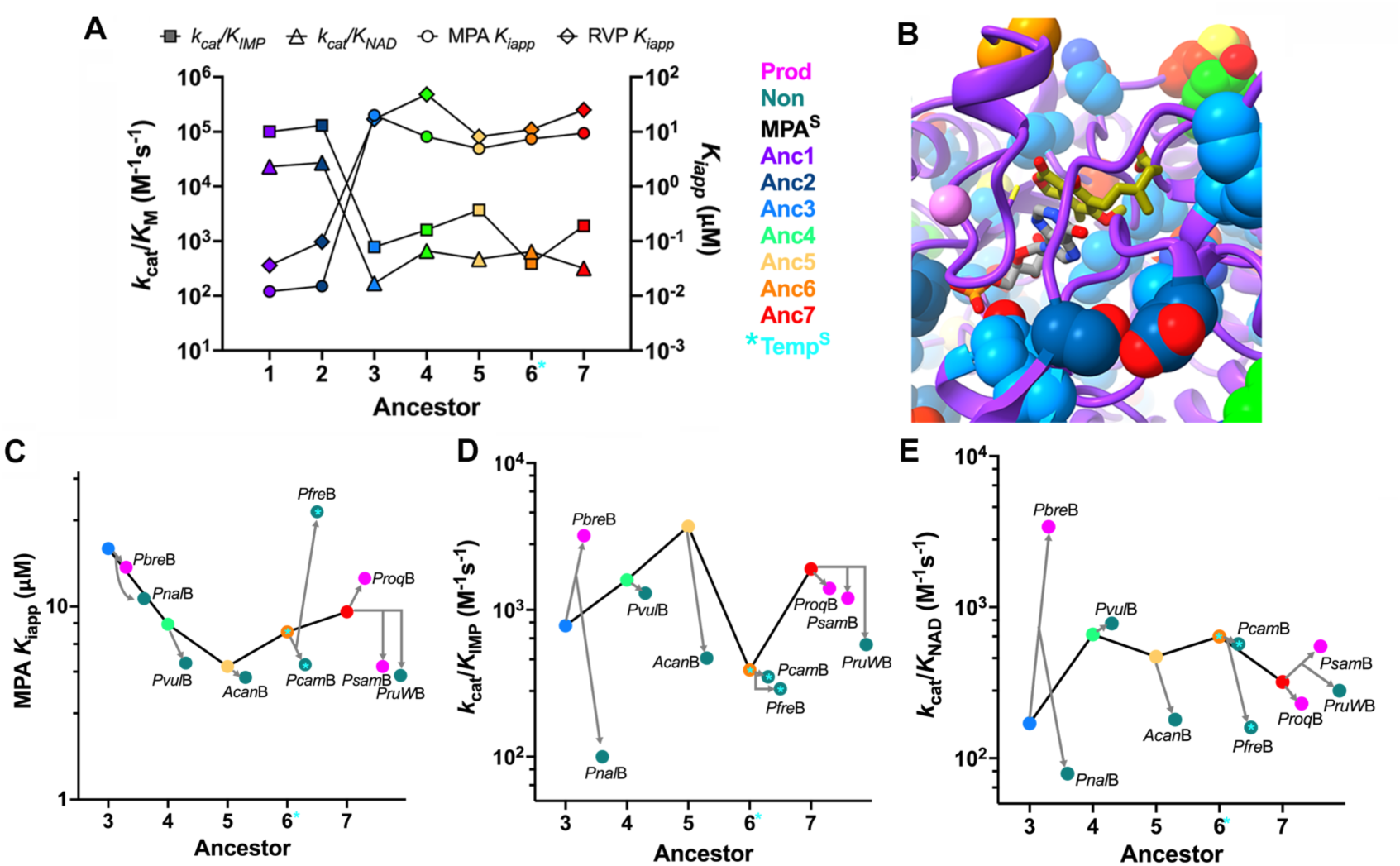
Evolution of IMPDH-Bs. **A.** Timelines for the changes in resistance (values of *K_iapp_* for MPA and RVP) and catalytic efficiency (*k_cat_/K_IMP_* and *k_cat_/K_NAD_*) for the resurrected ancestors. **B.** Ancestor substitutions near the MPA binding site. The structure of Anc1 (purple) was predicted using AlphaFold2 (Figure S3; (81)). Substitutions in Anc2-7 are shown colored as in the key in panel **A**. K^+^, pink ball; IMP, gray sticks; MPA, olive sticks. These ligands were positioned by aligning Anc1 with the *C. griseus* E-XMP*•MPA complex (PDB: 1JR1, (78)). **C-E.** Timelines for the changes in resistance (values of *K_iapp_* for MPA and RVP) and catalytic efficiency (*k_cat_/K_IMP_* and *k_cat_/K_NAD_*) for the ancestors and their extant descendants. MPA producers are magenta, nonproducers are teal, ancestors are colored as in the key in panel **A**.

### Resistance declines moderately after Anc3

Anc4-7 contained 10, 4, 5 and 11 substitutions relative to their respective predecessors (Figure S6). All of these enzymes are MPA-resistant with high values of *K_IMP_* and low values of *k_cat_/K_IMP_* and *k_cat_/K_NAD_* characteristic of IMPDH-Bs (Figures 2-G and 3A). Anc3 is more MPA-resistant than all of its descendants with the exception of *Pfre*B (Figure 3C). Even *Pbre*B, which must withstand industrial MPA levels that can reach 6g/L (52), is less resistant than Anc3. With the exceptions of *Pfre*B and *Proq*B, the descendants are all less resistant than their nearest ancestors, albeit still ∼100-fold more resistant than a typical eukaryotic IMPDH. It seems likely that resistance remained above the threshold needed to protect an organism from toxicity in all cases.

### Activity increases after Anc3, but declines in modern IMPDH-Bs

With the exception of Anc6, catalytic efficiency underwent further optimization after Anc3, increasing by 4 to 5-fold as measured by both *k_cat_/K_IMP_* and *k_cat_/K_NAD_* (Figure 3A,D,E). This initial period of activity optimization was followed by downward drift in the modern descendants. All modern IMPDH-Bs are less active than their nearest ancestor. These differences are more pronounced for modern enzymes from non-producers. *Pbre*B is the notable exception; values of *k_cat_/K_IMP_* and *k_cat_/K_NAD_* for *Pbre*B are 4- and 20-fold higher, respectively, than those of Anc3. The continued optimization of activity in *Pbre*B may reflect the added selective pressure of the industrial production of MPA.

### A resistance-activity tradeoff

Linear free energy relationships were observed between MPA inhibition and activity as measured by both *k_cat_/K_IMP_* and *k_cat_/K_NAD_* (Figure 4A,B). Higher catalytic efficiency correlated strongly with higher inhibition (lower resistance). In addition to IMPDH-Bs, ancestral and MPA^S^ enzymes, these correlations included 3 moderately MPA resistant IMPDHs (MPA^M^), *Pbre*A and the enzymes from *Cryptococcus neoformans* and *C. gatti* (53) for a total of 28 enzymes (Tables S1 and S2). The correlations span approximately 3 orders of magnitude, with a stronger dependence on the value of *k_cat_/K_IMP_* than *k_cat_/K_NAD_* (slopes = -0.65 ± 0.07 and - 0.44 ± 0.09 for *k_cat_/K_IMP_* and *k_cat_/K_NAD_*, respectively). These relationships reveal a strong activity-resistance tradeoff that persists throughout the eukaryotic IMPDH lineage.

**Figure 4.**
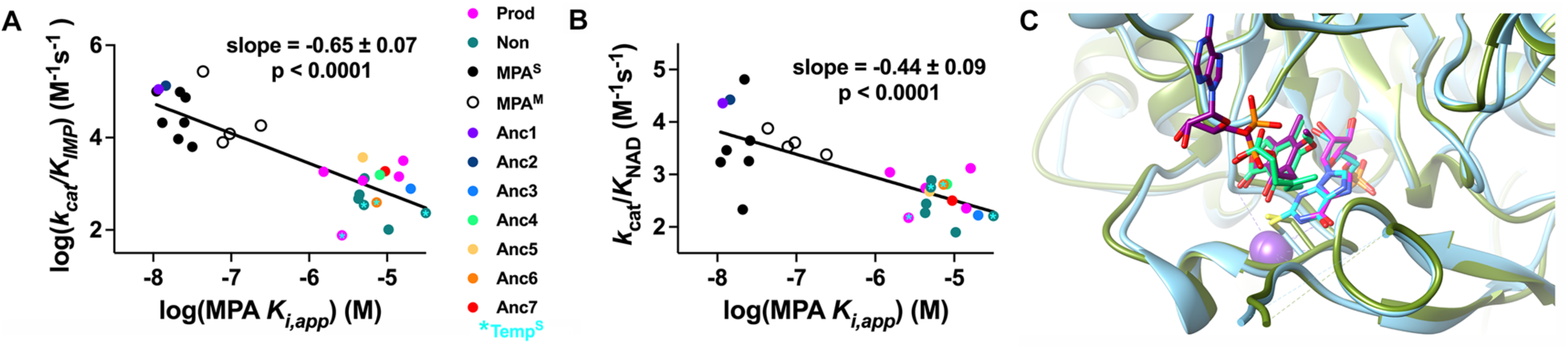
Tradeoffs between activity and resistance. **A.** Linear free energy relationship between the values of *k_cat_/K_IMP_* and MPA resistance (*K*_iapp_). MPA^S^, *K_iapp_* ≤ 30 nM; MPA-moderately resistant (MPA^M^), 1 μM ≥ *K_iapp_* > 30 nM; MPA^R^, *K_iapp_* > 1 μM. See Tables S1 and S2 for all values. **B.** Linear free energy relationship between *k_cat_/K_NAD_* and MPA resistance (*K*_iapp_). Key as in panel **A**. **C.** E•RVP mimics E-XMP*. The structures of E-XMP*•MPA (cornflower blue, PDB: 1JR1) (78) and E•RVP•MAD (olive, PDB: 1NF7; MAD = mycophenolate adenine dinucleotide) are overlaid. E-XMP* is shown in cyan, MPA in spring green, RVP in magenta, and MAD in dark magenta. Figure rendered in UCSF Chimera (79).

### Ancestral and modern IMPDH-Bs are also resistant to RVP

The IMP competitive inhibitors RVP, MZP and OxMP induce enzyme conformations that mimic E-XMP*, as illustrated in the overlay of the E-XMP*•MPA and E•RVP•MAD complexes (Figure 4C) (54–56). To determine if the activity-resistance tradeoff extended to IMP competitive inhibitors, we assayed RVP inhibition at 0.5x*K_IMP_* and 1.8x*K_NAD_* (Figure S8). As previously reported (57), RVP is a potent inhibitor of the MPA^S^ hIMPDH2, with only 15% activity observed under these conditions. The value of *K_i_* was estimated to be 0.12 µM assuming competitive inhibition, in reasonable agreement with the literature (0.39 µM (57)). *A. nidulans* IMPDH-A (*An*IMPDH-A) and Anc1 were also very sensitive to RVP (*K*_i_ ≤ 40 nM), while Anc2 was less sensitive by a factor of 2.7 (*K*_i_ = 96 nM) (Tables S1 and S2). Remarkably, most ancestral and modern IMPDH-Bs had >90% activity in the presence of RVP (Figure S8), indicating values of *K_i_* ≥ 40 µM. *Apse*B, *Proq*B and *PruW*B were exceptions, with activity ranging from 57-89%, corresponding to *K_i_* values of 0.9, 5.2 and 0.97 µM, respectively. Two IMPDH-As, *Pbre*A and *PruW*A, were also moderately resistant to RVP (*K_i_* = 3.0 µM and 0.92 µM, respectively).

## Discussion

### The evolution of self-resistance

Gene duplications/amplifications are pivotal events in the evolution of new enzyme functions (58–61). These events are especially critical in the development of drug resistance because increased gene dosage alone often creates a transient resistant state that permits the emergence of beneficial point mutations (58–60, 62, 63). Gene amplification has often been observed in laboratory selections for MPA resistance (39–42), so it is reasonable to posit that the initial duplication event leading to IMPDH-Bs was a response to the presence of MPA or another IMPDH inhibitor. Interestingly, while Anc2 is MPA^S^ (Table S2), it displays 2.7-fold higher resistance to RVP than Anc1. This observation suggests that the duplication could have been a response to a similar IMP-competitive inhibitor, creating a genetic background amenable to the production of MPA.

MPA resistance emerged between Anc2 and Anc3, accompanied by a decrease in enzyme activity (Figure 3A). With one exception, Anc3 was more MPA resistant than any of its descendants. Even *Pbre*B was less resistant than Anc3, although *Pbre*B must be active during the industrial production of MPA, where concentrations can reach 6 g/L (52). The burst of adaptive evolution that created Anc3 appears to have exceeded the threshold required for MPA production, allowing subsequent genetic drift to erode resistance. Similar bursts of innovation followed by downward functional drift have been noted in other studies of enzyme evolution (64). Alternatively, the decline in resistance may have been necessary to permit the optimization of enzyme activity.

### Why do non-producers retain IMPDH-Bs?

Our observations are consistent with the hypothesis that MPA biosynthesis emerged prior to the radiation of *Pencillium*, with subsequent BGC loss in many lineages (30, 45). Yet while the BGC appears to be dispensable, functional, IMPDH-Bs appear to be preserved in most *Penicillium,* albeit somewhat less active. Duplicate genes typically decline to nonfunctionalization in the absence of purifying selection (65, 66), suggesting *mpaF* must confer a fitness advantage. MPA is a common environmental mycotoxin (8), so perhaps MPA is encountered with sufficient frequency to preserve *mpaF* in nonproducers. Although little is known about the prevalence of mizoribine and oxanosine in the environment, these IMP-competitive inhibitors are also produced by soil microbes that may co-exist with MPA nonproducers. Therefore *mpaF* may be retained in MPA nonproducers to protect against IMPDH inhibitors in the environment.

### The complicated evolution of IMPDH-B and the MPA BGC

The incongruence of the IMPDH-B phylogeny and the fungal species trees suggests a complicated evolutionary history likely involving both vertical and horizontal gene transfer. Given the propensity of MPA to induce gene amplification, incomplete lineage sorting is also a possibility (67). Although the relatively small number of extant BGCs makes the evolution of the MPA BGC difficult to assess, it seems likely that the cluster experienced a similarly complex evolutionary trajectory, as evidenced by the sporadic presence of BGC throughout the species tree. Our discovery of a likely horizontal gene transfer between *Aspergillus* and *B. moriformis* implies that BGC inheritance was not strictly vertical, in contrast to the hypothesis put forward in a recent 14-sequence phylogeny of *mpaC* (45). Intriguingly, based on enzyme activity, the Anc4, 5 and 7 fungi appear to be MPA producers while Anc6 fungus was likely a nonproducer, suggesting that the BGC may have been silenced or lost and subsequently reactivated or reacquired. Perhaps the Anc6 fungus or a predecessor migrated to a colder environment where MPA production was not beneficial. Phylogenetic investigations of the other genes in the MPA BGC may be able to determine if the BGC has been lost and required.

### A design-constraint tradeoff

Tradeoffs are commonly believed to shape the evolution of new enzyme functions (68). Stability-activity tradeoffs are a common paradigm (69). A mutation that changes the architecture of the active site typically destabilizes the structure, but stability can usually be restored with compensating mutations elsewhere in the protein (70). Thus while stability-activity tradeoffs can profoundly shape evolutionary trajectories, these two traits are largely independent. The tradeoff is probabilistic, arising because the chances that a function-altering mutation will stabilize structure are low (71). Given enough time, a stable active enzyme will emerge (72, 73).

In contrast, the experiments described above reveal a strong tradeoff between activity and resistance that has likely persisted for many millions of years. Shankovich and colleagues have reported a similar activity-resistance tradeoff in a 20 day directed evolution experiment producing trimethoprim-resistant dihydrofolate reductase (DHFR) (74, 75). The magnitude of this tradeoff was more modest than that observed in our experiments: the slope of the log(*k_cat_/K_m_*) versus log(*K_i_*) plot was -0.3 as compared to -0.65 with IMPDH activity and MPA resistance (Figures 4A and S9). Importantly, active resistant DHFRs are observed in clinical strains, where trimethoprim resistance can increase 500-fold without an accompanying activity penalty (76). Therefore it seems likely that the activity-resistance tradeoff observed in Shankovich’s experiment would resolve in a longer selection.

We propose that the activity-resistance tradeoff in fungal IMPDHs derives from a design restraint rather than a probabilistic constraint. Selections for MPA-resistant eukaryotic IMPDHs have yielded enzymes that are at best modestly resistant (4-fold) (33, 34). These observations suggest that catalytically active, highly MPA-resistant IMPDHs are not readily accessible in the eukaryotic IMPDH lineage, so acquisition of resistance must be accompanied by a loss of enzyme efficiency. The apparent inability to resolve the activity-MPA resistance conflict is also suggested by the observation that all fungi containing IMPDH-Bs, with the exception of *P. parvum,* also contain IMPDH-A, suggesting that IMPDH-Bs are not able to support growth on their own.

Activity and MPA resistance nevertheless can be uncoupled, as demonstrated by prokaryotic IMPDHs that have both high activity and ∼10^3^-fold resistance to MPA relative to sensitive to eukaryotic enzymes (25). Prokaryotic IMPDHs are highly diverged from eukaryotic enzymes, as dramatically illustrated by the relocalization of the cofactor binding site (77). Therefore it is quite possible that epistasis blocks the evolutionary trajectories to highly active MPA-resistant enzymes like prokaryotic IMPDHs. However, unlike IMPDH-Bs, prokaryotic IMPDHs are sensitive to RVP and similar IMP competitive inhibitors (25). This observation suggests a more intriguing explanation: fungi may require resistance to both MPA and IMP-competitive inhibitors like RVP. Modest MPA resistance can result from substitutions that stabilize the closed conformation of E-XMP*, which blocks MPA access (34). However, such substitutions also cause hypersensitivity to IMP-competitive inhibitors (34). MPA binds to E-XMP*, while RVP, OxMP and MZP form complexes that closely resemble E-XMP* (54–56, 78). Therefore the inhibitors bind to an enzyme conformation required for catalysis, creating a design constraint that produces the activity-resistance tradeoff.

## Acknowledgements

This work was funded in part by National Institute of Health grants R01AI125362 (L.H.) and T32EB009419 (A.W.S.), Novo Nordisk Foundation grant NNF18OC0034952 (J.L.S.) and Danish National Research Foundation DNRF137 (J.C.F.)

## Supporting Material

**Figure S1.**
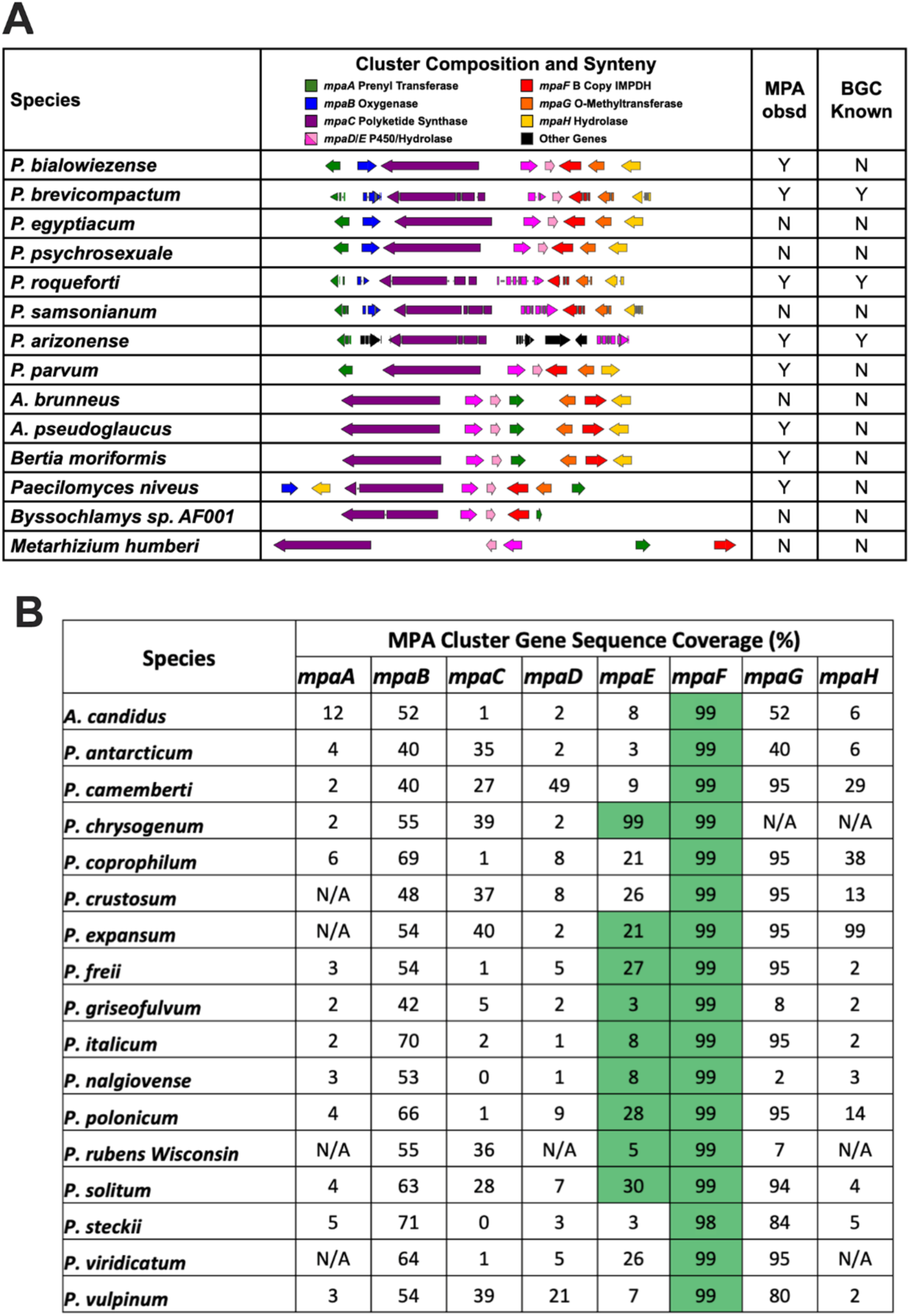
MPA biosynthetic gene clusters in filamentous fungi. **A.** Organization of MPA BGC in producers. References for previously reported BCGs: *A. brunneus* (1), *A. pseudoglaucus* (2), *Paecilomyces niveus* (*Pani*) (3), *P. arizonense* (*Pari*) (4, 5), *P. brevicompactum* (6), *P. bialowiezense* (7), *P. parvum* (8), and *P. roqueforti* (9). Some clusters annotate *mpaD* and *mpaE* as a combined gene; this is depicted by only showing the darker pink color. **B.** Many MPA nonproducers possess remnants of MPA BGC on the same genomic contig as *mpaF*. All *mpaF* sequences have ≥98% sequence coverage relative to *P. brevicompactum mpaF.* Highlighted in green are the BLAST matches with sequence coverage from ≥8 to 100% and sequence identity ≥65% on the same contig as *mpaF*.

**Figure S2.**
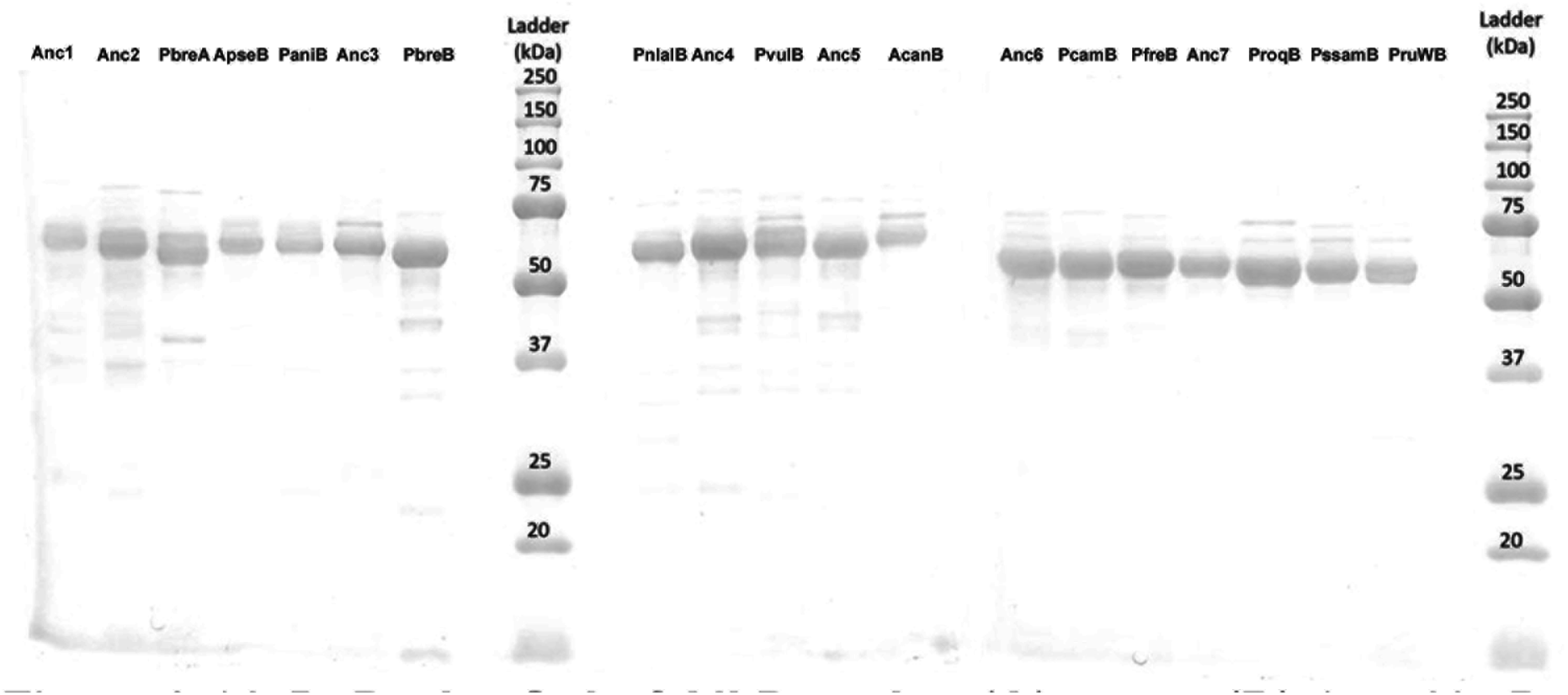
Purification of fungal IMPDHs. Proteins (1 μg) were resolved on SDS-PAGE (12% acrylamide) and stained with Coomassie brilliant blue.

**Figure S3.**
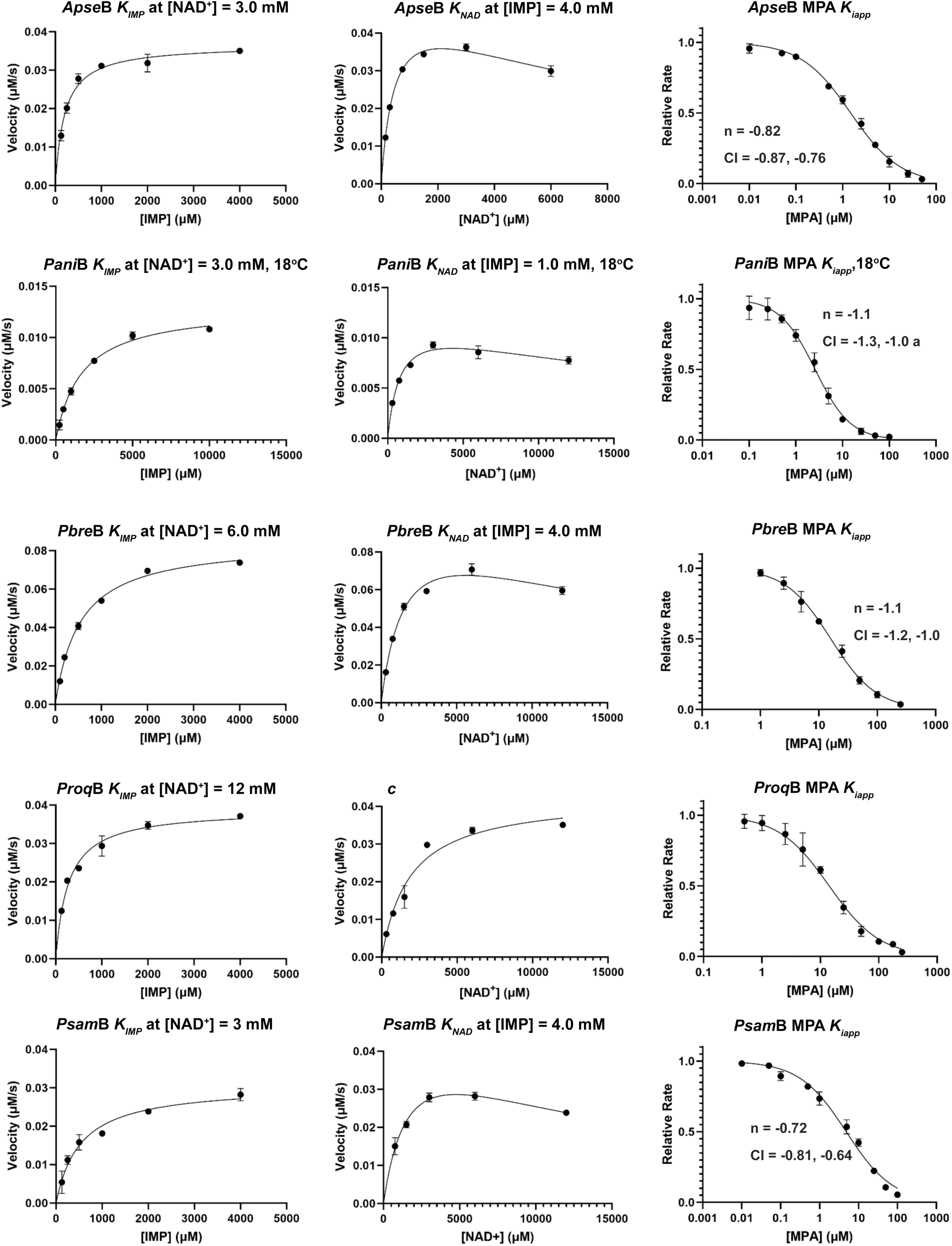
Kinetic characterization of producer IMPDH-Bs. Enzyme concentrations are listed in the Methods section. Reactions were with fixed substrate concentrations as listed and at 25°C unless otherwise specified. The average and range of duplicate reactions are plotted. Data was analyzed in GraphPad Prism. The values of *K_IMP_* were determined by fits to the Michaelis-Menten equation, values of *K_NAD_* were determined by fits to Michealis-Menten or substrate inhibition equation as appropriate. The values of *K_iapp_* for MPA were determined as described in the Methods. CI, 95% confidence intervals. Values are listed in Table S1.

**Figure S4.**
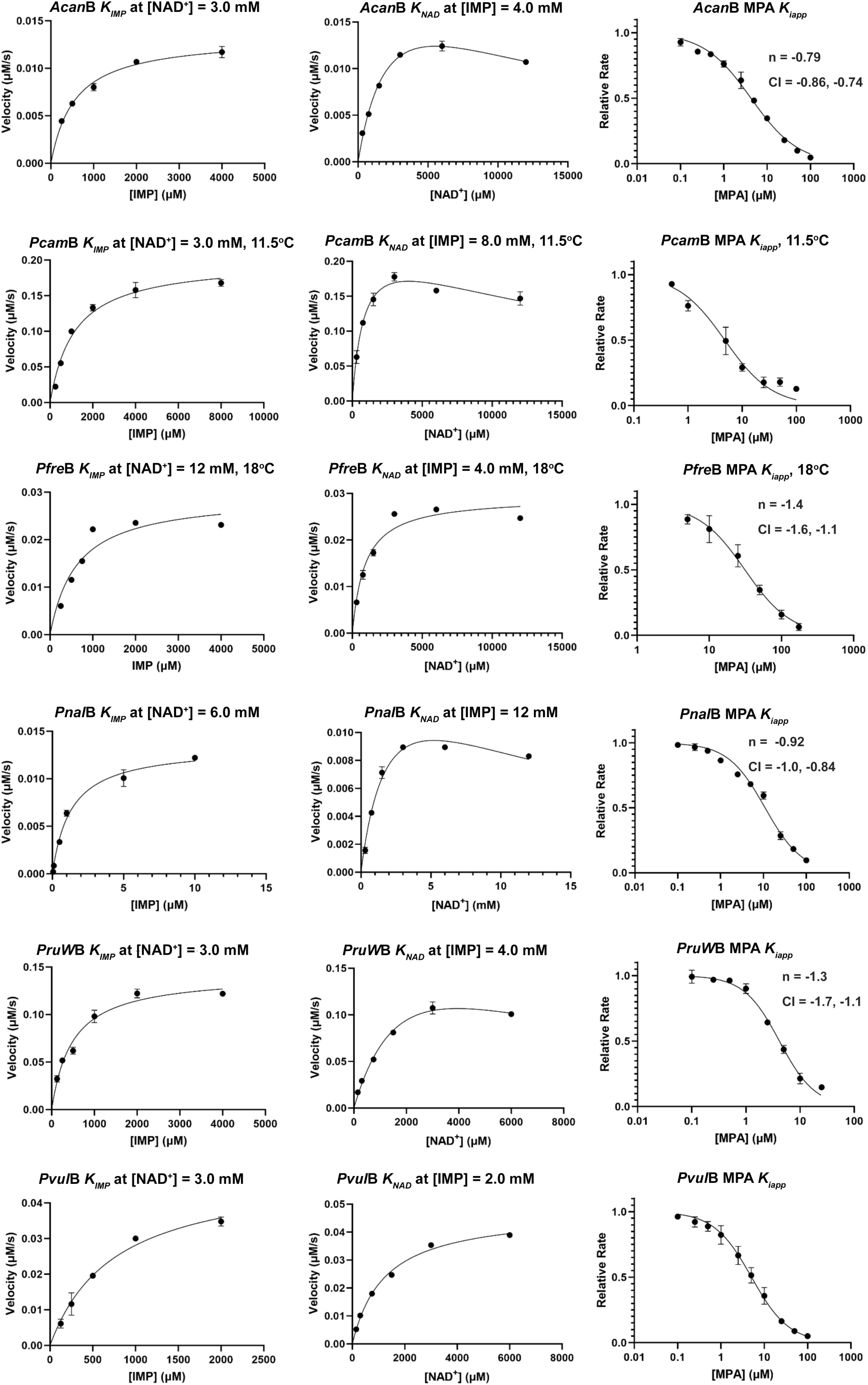
Kinetic characterization of non-producer IMPDH-Bs. Enzyme concentrations are listed in the Methods section. Reactions were with fixed substrate concentrations as listed and at 25°C unless otherwise specified. The average and range of duplicate reactions are plotted. Data was analyzed in GraphPad Prism. The values of *K_IMP_* were determined by fits to the Michaelis-Menten equation, values of *K_NAD_* were determined by fits to Michealis-Menten or substrate inhibition equation as appropriate. The values of *K_iapp_* for MPA were determined as described in the Methods. CI, 95% confidence intervals. Values are listed in Table S1.

**Figure S5.**
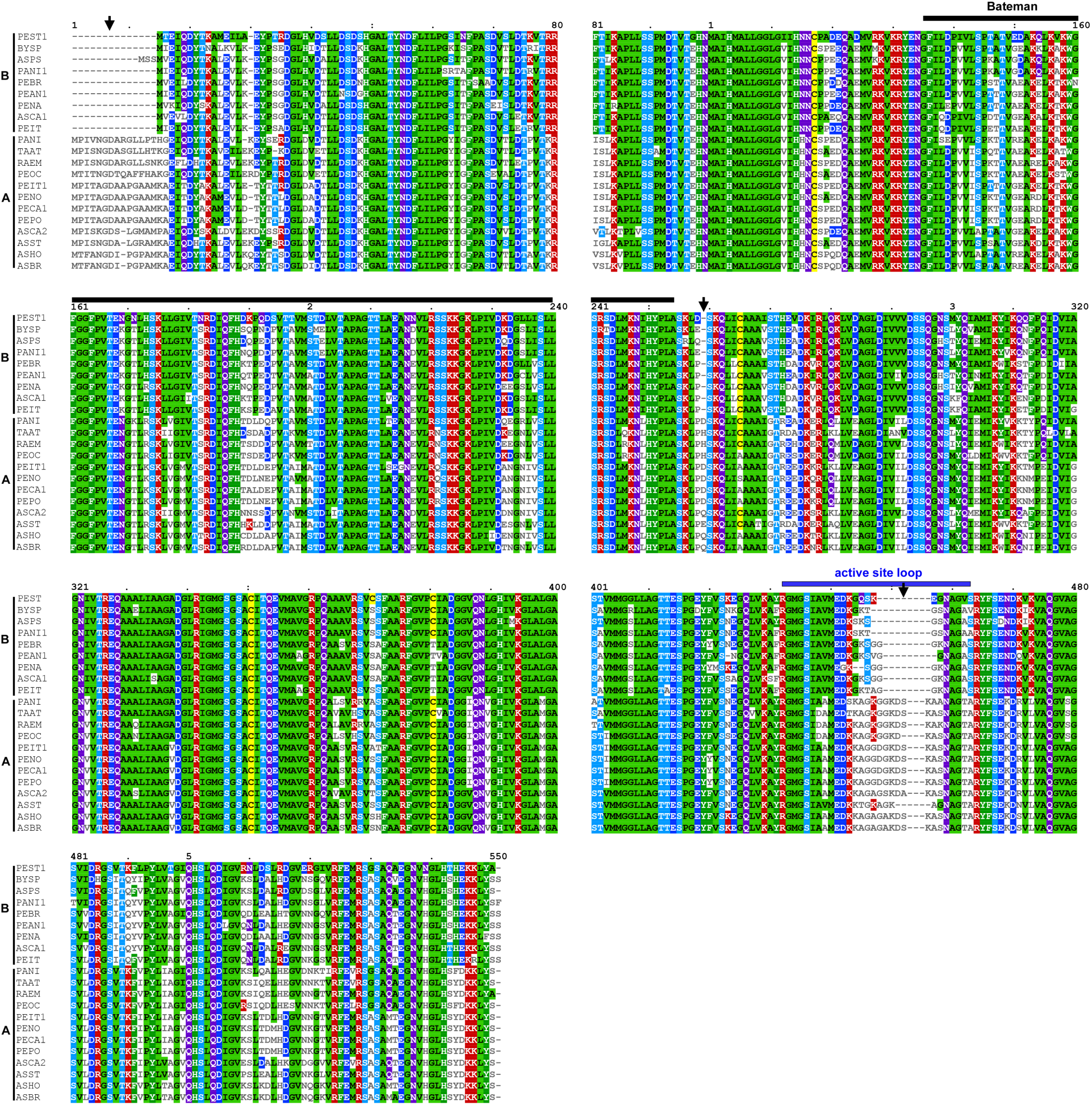
Sequence alignment of IMPDH-As and -Bs. Arrows mark the deletions that are characteristic of IMPDHBs. Representative sequences are shown. The full 83 sequence alignment can be downloaded at https://github.com/alexandersarkis/EvolutionOfDrugResistanceInPenicilliumFungiAWS.

**Figure S6.**
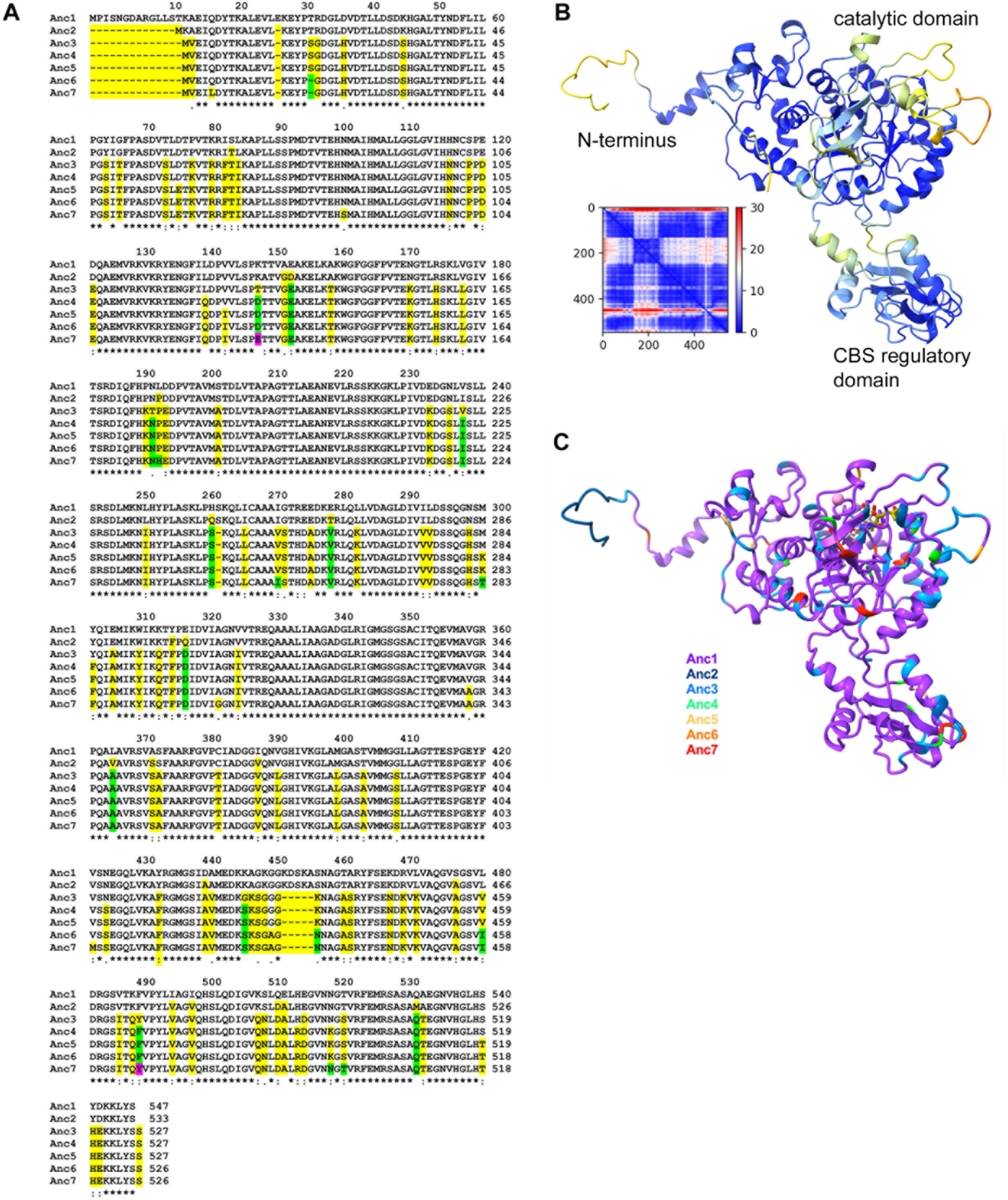
Ancestral IMPDHs. **A**. Clustal Omega sequence alignment of ancestral IMPDHs (10). Ancestors listed in order of appearance in the tree in Figure 2A. First change at a given position denoted in yellow, second change in green, third change in magenta. **B.** AlphaFold model of Anc1 colored by pLDDT (11). Inset shows PAE. **C.** Anc1 is shown in purple ribbon, substitutions in Anc2-7 are shown colored as in the key. K^+^, pink ball. IMP, gray sticks. MPA, olive sticks. These ligands were positioned by aligning Anc1 with the *C. griseus* E-XMP*•MPA complex (PDB: 1JR1, (12)).

**Figure S7.**
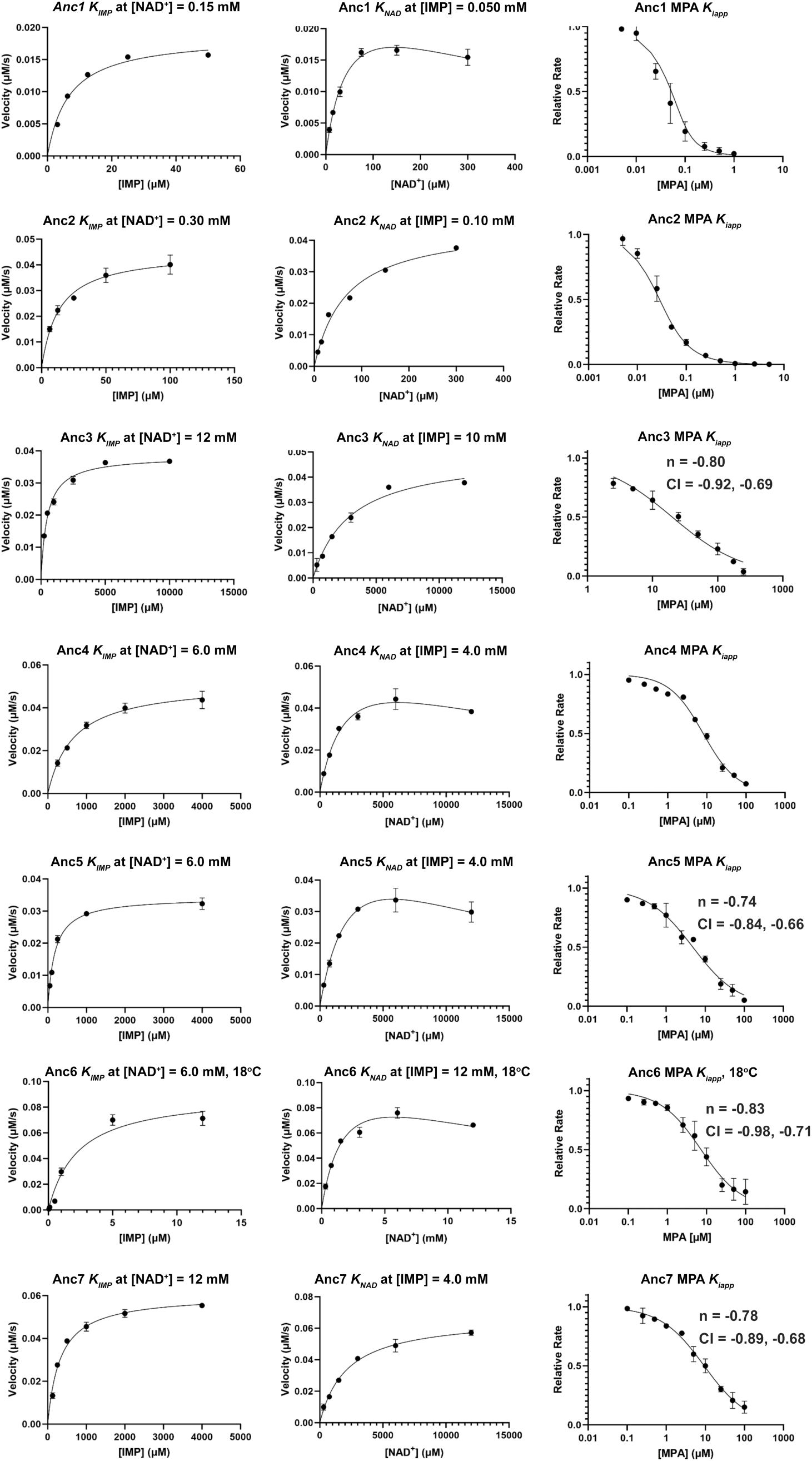
Kinetic characterization of IMPDH-B Ancestors. Enzyme concentrations are listed in the Methods section. Reactions were with fixed substrate concentrations as listed and at 25°C unless otherwise specified. The average and range of duplicate reactions are plotted. Data was analyzed in GraphPad Prism. The values of *K_IMP_* were determined by fits to the Michaelis-Menten equation, values of *K_NAD_* were determined by fits to Michealis-Menten or substrate inhibition equation as appropriate. The values of *K_iapp_* for MPA were determined as described in the Methods. CI, 95% confidence intervals. Values are listed in Table S1.

**Figure S8.**
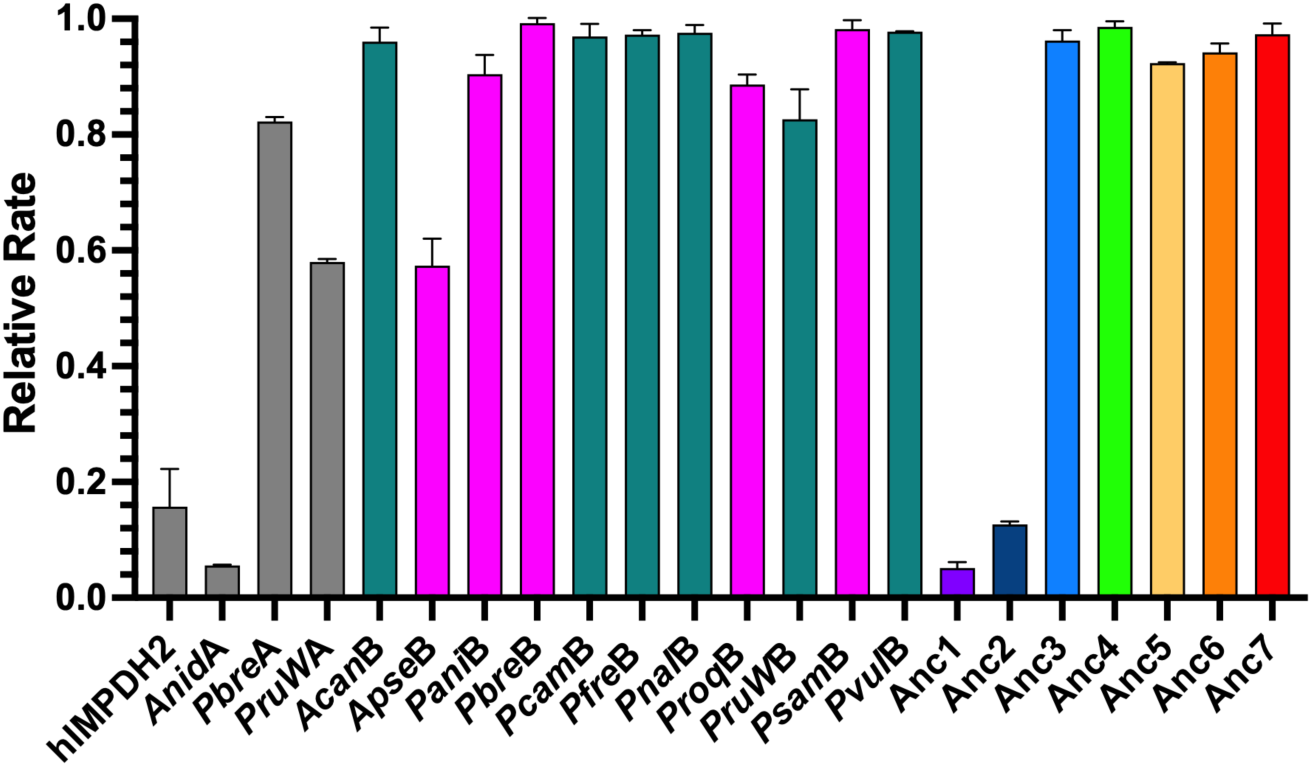
RVP inhibition of fungal IMPDHs. Enzyme concentrations are listed in Methods section. Reactions performed in duplicate at 0.5\**K_IMP_* and 1.8\**K_NAD_* and 1 mM RVP at 25°C unless otherwise noted. Anc6, *Pani*B and *Pfre*B were assayed at 18°C. *Pcam*B was assayed at 11.5°C. Gray denotes representative eukaryotic IMPDHs, magenta denotes enzymes from MPA producers, teal denotes enzymes from nonproducers and ancestors are colored as in Figures 2-4.

**Figure S9.**
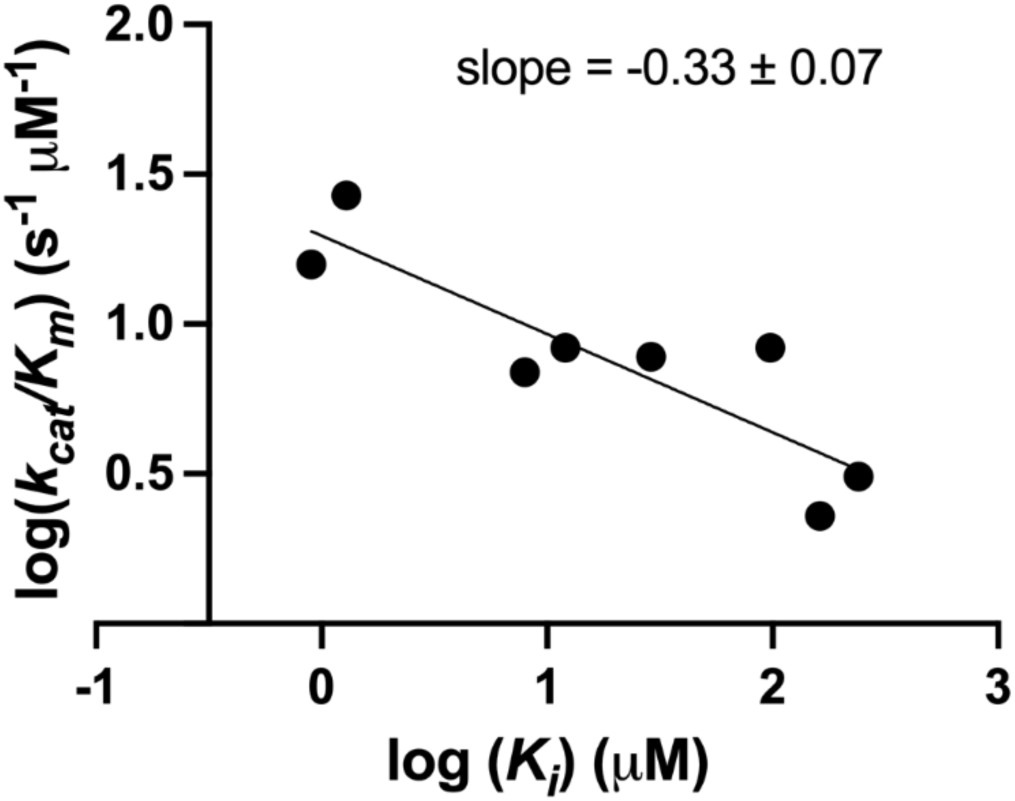
DHFR-trimethoprim activity-resistance tradeoff. Data from (13).

**Figure S10.**
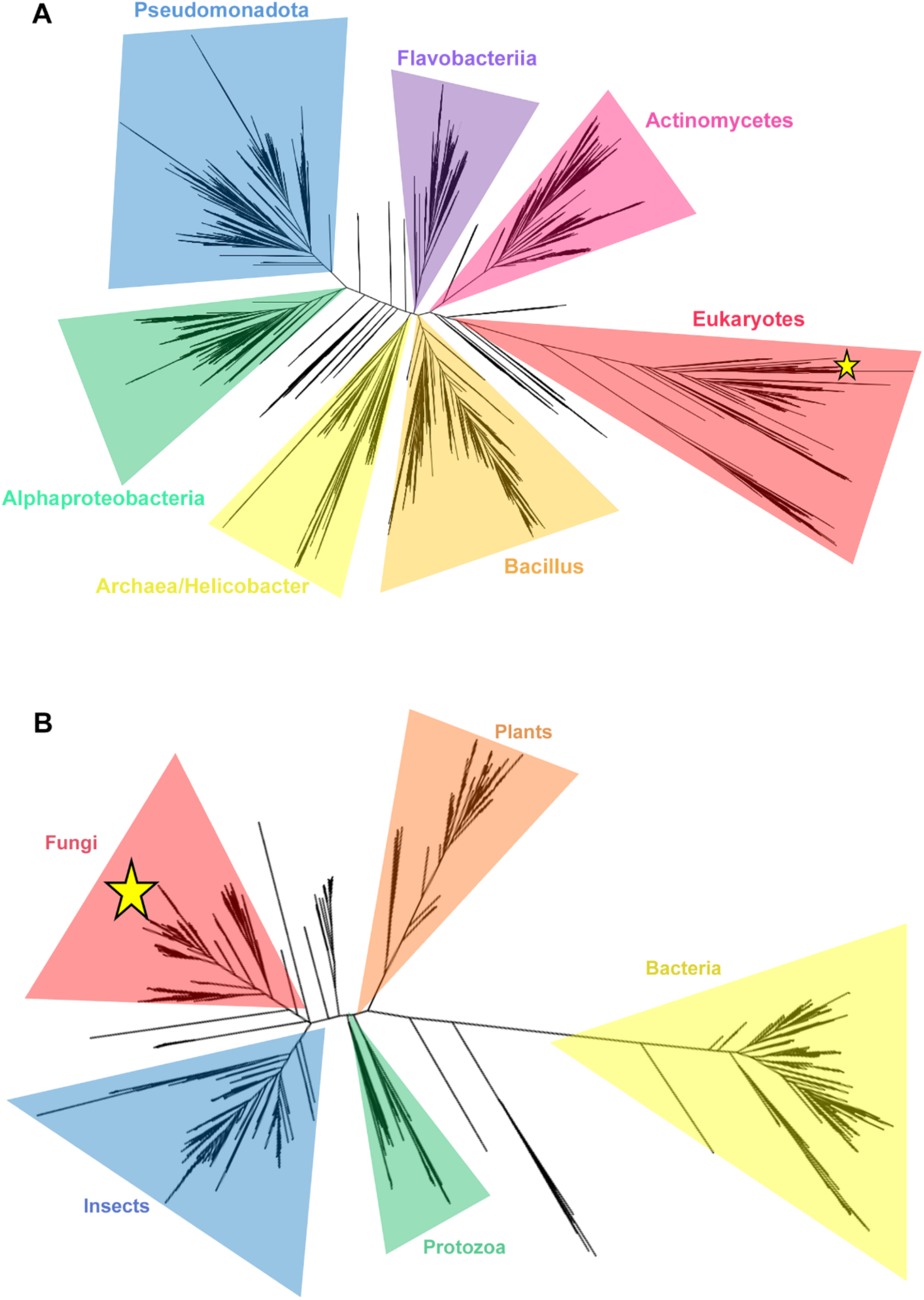
IMPDH phylogeny. A. Comprehensive phylogenetic tree of 20059 IMPDH sequences consisting mainly of bacterial clades. B. Smaller tree focused on 1371 eukaryotic sequences, with bacterial IMPDHs as the outgroup. The approximate location of *Pbre*B is denoted with a star. Alignments can be downloaded at https://github.com/alexandersarkis/EvolutionOfDrugResistanceInPenicilliumFungiAWS

**Table S1.**
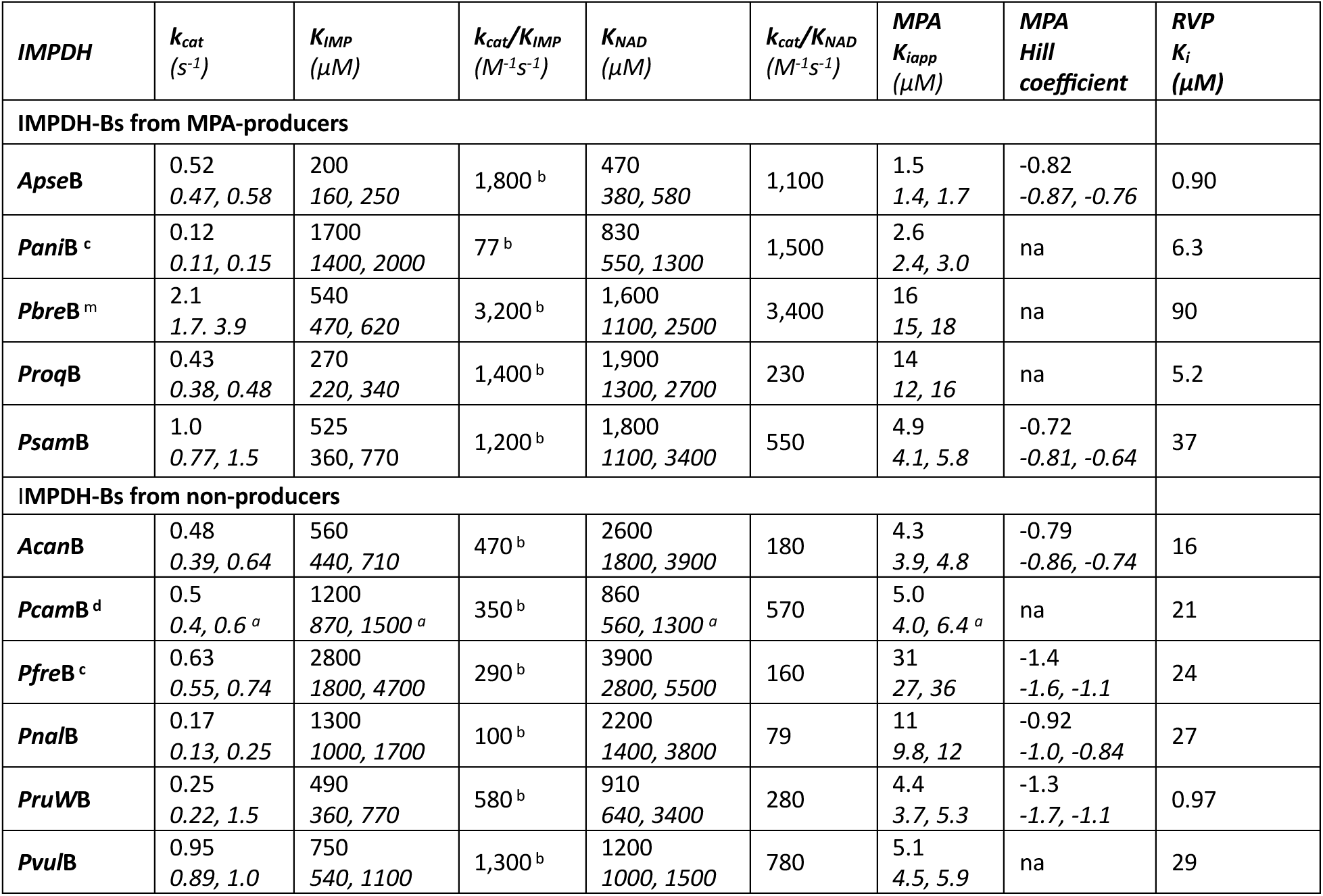
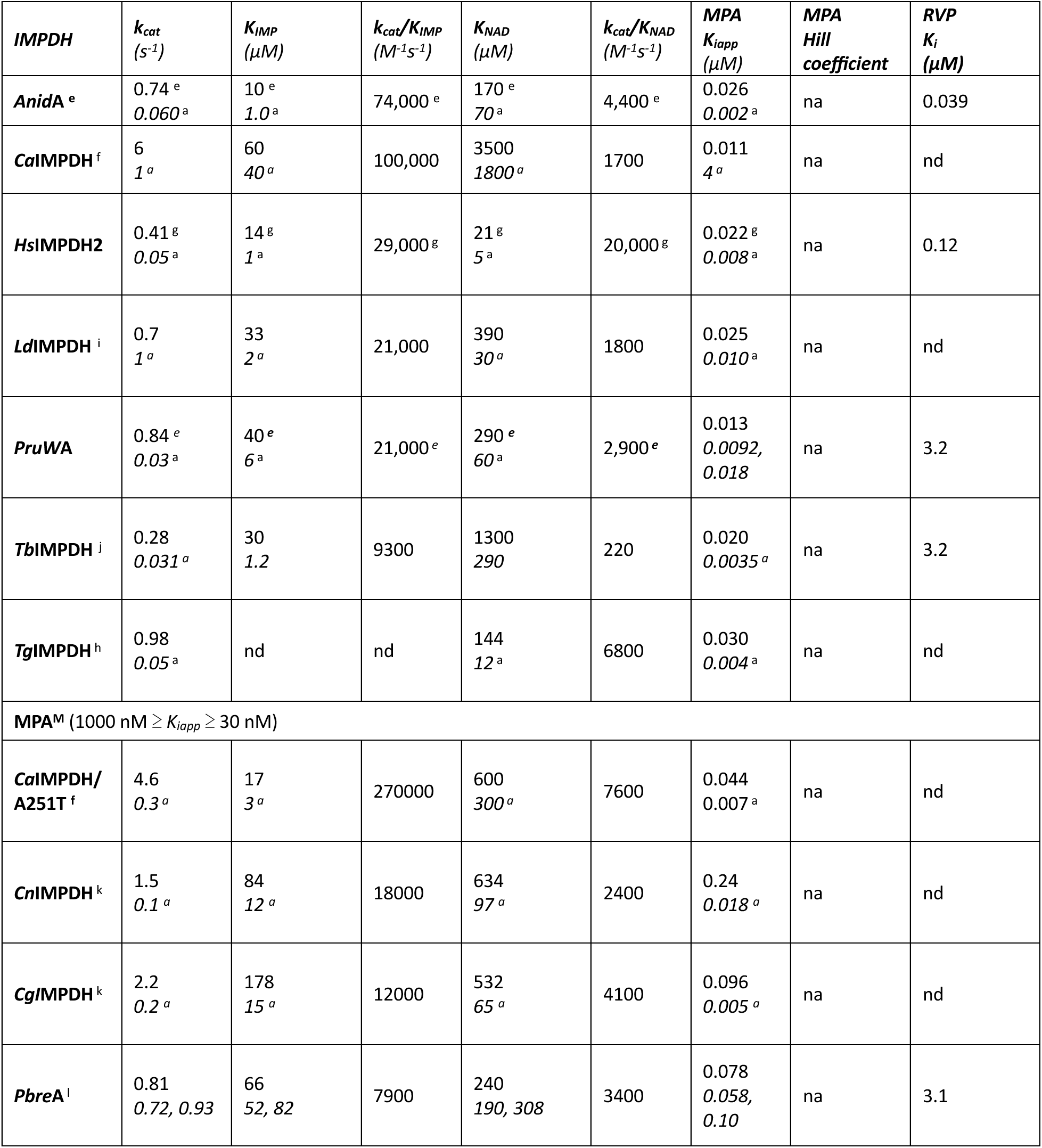
Characterization of modern IMPDHs. Assays were performed in 50 mM Tris-HCl, pH 8, 100 mM KCl and 1 mM DTT at 25°C unless otherwise noted. Italics denote 95% confidence limits; na, not applicable; nd, no data. MPA^S^, MPA-sensiKve; MPA^M^, moderately MPA resistant. *Acan, Aspergillus candidus; Anid, A. nidulans, Apse, A. pseudoglaucus*; *Ca, Candida albicans; Cg, Cryptococcus gattii; Cn, C. neoformans; Hs, Homo sapiens; Ld, Leishmania donovani; Pani, Paecilomyces niveus*; *Pbrev, Penicllium brevicompactum; Pbia, P. bialowiezense*; *Pcam*, *P. camemberA*; *Pfre*, *P. freii*; *Pnal*, *P. nalgiovense*; *Ppar*, *P. parvum; Proq, P. roqueforA*; *PruW*, *P. rubens Wisconsin; Psam, P. samsonianum; Pvul, P. vulpinum; Tb, Trypanosoma bruceii; Tg, Toxoplasma gondii;* a. Italics denote standard deviaKon. b. Sub-saturaKng NAD^+^ condiKons at or above *KNAD* were used due to NAD^+^ substrate inhibiKon. c. Assays performed at 18°C. d. Assays performed at 11.5°C. e. Values from (14). f. Values from (15). g. Values from (16). h. Values from (17). i. Values from (18). j. Values from (19). k. Values from (20). l. *Pbre*A literature values: *kcat* = 0.70 s^−1^, *KIM*P = 130 μM, K*NAD* = 340 μM (14); *kcat* = 2.7 s^−1^, *KIM*P = 93 μM, K*NAD* = 400 μM. m. *Pbre*B literature values: *kcat* = 0.41 s^−1^, *KIM*P = 1400 μM, K*NAD* = 790 μM (14);

**Table S2.** Characterization of ancestral IMPDHs. Assays were performed in 50 mM Tris-HCl, pH 8, 100 mM KCl and 1 mM DTT at 25°C unless otherwise noted. Italics denote 95% confidence limits. na, not applicable. a. Sub-saturating NAD^+^ conditions at or above *KNAD* were used due to NAD^+^ substrate inhibition. b. Assays performed at 18°C.

| **IMPDH** | **$k_{cat}$**<br>**($s^{-1}$)** | **$K_{IMP}$**<br>**($\mu M$)** | **$k_{cat}/K_{IMP}^a$**<br>**($M^{-1}s^{-1}$)** | **$K_{NAD}$**<br>**($\mu M$)** | **$k_{cat}/K_{NAD}$**<br>**($M^{-1}s^{-1}$)** | **MPA $K_{iapp}$**<br>**($\mu M$)** | **RVP $K_i$**<br>**($\mu M$)** |
| --- | --- | --- | --- | --- | --- | --- | --- |
| **Anc1** | 1.2<br>*0.91, 1.6* | 6.7<br>*5, 8.9* | $1.10 \times 10^5$ | 50<br>*33, 84* | 23,000 | 0.012<br>*0.0018, 0.028* | 0.036<br>*0.022, 0.050* |
| **Anc2** | 1.8<br>*1.6, 2* | 14<br>*9.5, 19* | $1.30 \times 10^5$ | 67<br>*51, 90* | 27,000 | 0.015<br>*0.0092, 0.021* | 0.097<br>*0.090, 0.10* |
| **Anc3** | 0.49<br>*0.44, 0.56* | 490<br>*400, 600* | 780 | 3,000<br>*2200, 4000* | 170 | 20<br>*17, 24* | 17<br>*17, 17* |
| **Anc4** | 1.5<br>*1.1, 2.4* | 670<br>*500, 880* | 1,600 | 2,300<br>*1300, 4600* | 660 | 8.1<br>*7.2, 9.2* | 48<br>*48, 48* |
| **Anc5** | 1.3<br>*0.97, 2.3* | 180<br>*150, 220* | 3,700 | 2,900<br>*1600, 6100* | 470 | 4.9<br>*4.1, 5.8* | 8.1<br>*8.1, 8.1* |
| **Anc6<sup>b</sup>** | 1.2<br>*0.87, 2.0* | 2,300<br>*1400, 4000* | 390 | 1,900<br>*980, 4200* | 640 | 7.4<br>*6.0, 9.0* | 11<br>*11, 11* |
| **Anc7** | 0.67<br>*0.63, 0.72* | 320<br>*260, 390* | 1,900 | 2,100<br>*1700, 2600* | 320 | 9.4<br>*7.9, 11* | 25<br>*25, 25* |

## MATERIALS AND METHODS

### Materials

NAD^+^ was purchased from ThermoFisher, IMP from MP Biomedicals and MPA from Sigma.

### IMPDH Purification

Expression systems for *Pbre*B, *Pbre*A, *An*ImdA and *Pru*WB (formerly named *P. chrysogenum* IMPDH-B) were reported previously (14). All other IMPDH sequences were codon optimized for *E. coli* expression and cloned into pET-28a(+) vectors (Novagen). Enzymes were expressed with N-terminal 6His tags in BL21(DE3)Δ*guaB* bacteria (21). One liter of Expression Medium (LB medium supplemented with 500 mM NaCl and 0.5% (v/v) glycerol) was inoculated with 50 mL of overnight culture grown in LB with 50 μg/ml kanamycin. Cells were grown at 37°C to an OD_600_ of 0.8-1.8 after which 0.5 mM IPTG was added to induce expression and cultures were shifted to 18°C overnight. Cells were harvested by centrifugation at 5000 g for 20 min at 4°C. The pellet was transferred into a 50 mL conical tube and resuspended in Lysis Buffer (50 mM K_2_HPO_4_, pH 8.0, 500 mM KCl, 5 mM imidazole, 10% glycerol (v/v) and 0.2 mM TCEP). IMP was added to a final concentration of 4 mM and cells were lysed by sonication. The lysate was clarified by centrifugation at 16,000 g for 30 min at 4°C and the supernatant was filtered through a 0.4 μm filter followed by a 0.22 μm filter. The lysate was then applied to a 3-5 mL gravity column of Super Ni NTA Resin (Anatrace) equilibrated with Lysis Buffer. The column was washed with 100 mL of Wash Buffer (Lysis Buffer with imidazole concentration increased to 25 mM) and enzyme was eluted with Elution Buffer (50 mM K_2_HPO_4_, pH 8.0, 100 mM KCl, 250 mM imidazole, 10% glycerol (v/v), 0.2 mM TCEP). IMPDH-containing fractions were identified by SDS-PAGE and activity assays and pooled. IMP (4 mM final) was added and the sample was dialyzed against 2 L of prechilled Dialysis Buffer 1 (50 mM K_2_HPO_4_, pH 8.0, 100 mM KCl, 3 mM EDTA, 1 mM TCEP, 10% glycerol (v/v)) followed by two transfers into Dialysis Buffer 2 (50 mM K_2_HPO_4_, pH 8.0, 100 mM KCl, 1 mM DTT, 10% glycerol (v/v)). Samples were concentrated in 30 kDa MilliporeSigma™ Amicon™ Ultra-15 Centrifugal Filter Units and aliquoted for storage. Protein concentraRon was determined in triplicate using the NanoOrange™ Protein Quantitation Kit (Fisher Scientific), using bovine serum albumin as the standard.

### IMPDH Activity Assays

IMPDH activity was monitored by measuring the increase in A_340_ from the formation of NADH at 25°C unless otherwise specified using a temperature controlled Shimadzu UV-2600 UV-Vis spectrophotometer. Assay Buffer contained 100 mM KCl, 50 mM Tris-Cl, pH 8.0, 3 mM EDTA, and 1 mM DTT with varied concentrations of IMP and NAD^+^. Initially, IMP was fixed at saturating concentrations while NAD^+^ concentration was varied. Substrate inhibition was often observed at high NAD^+^ concentrations, which prevented varying IMP concentrations at saturating NAD^+^. In this event, NAD^+^ was fixed at the concentration that gave the maximum rate, which was above the value of *K_NAD_* in all cases. Assays contained 50 nM enzyme with the exceptions of: 25 nM for Anc1, Anc2, and *PruW*A; 100 nM for Anc3, Anc6, Anc7, *Apse*B, *Pnal*B and *Proq*B; 200 nM for *Pfre*B; 500 nM for *Pcam*B, and *PruW*B. Kinetic parameters, *V_max_* and *K_m_*, were determined by the fit to the Micahelis-Menten equation (Equation 1) with GraphPad Prism, where *v* is the measured initial velocity and *S* is the concentration of varied substrate.

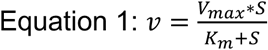

If NAD^+^ substrate inhibition was observed, initial rates were fit to Equation 2:

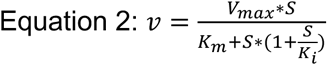

### MPA Inhibition Assays

Enzyme concentrations were as described above and IMP and NAD^+^ concentrations were fixed at 2 mM IMP and 1.5 mM, respectively. Assays were performed at 25°C unless otherwise noted. MPA concentrations were varied, and the assays performed in duplicate. The value of *K_iapp_* was calculated with GraphPad Prism using either Equation 3 or 4:

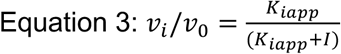

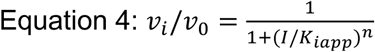

Where *v*_o_ is rate in the absence of inhibitor, *v*_i_ is the rate in the presence of inhibitor and *I* is the inhibitor concentration. In cases of tight binding inhibition (*I* ≈ *E*), *K_iapp_* was determined using the Dynafit (22) using the following code, where “I” is inhibitor concentration, and E is enzyme concentration:

~~~
[task]
 data = equilibrium
 task = fit
 approx = king-altman
 confidence = search
[settings]
 {ConfidenceIntervals}
  LevelPercent = 95
{Output}
 XAxisLabel = [MPA] (uM)
 YAxisLabel = Relative Rate
 BlackBackground = n
[mechanism]
 I + E <==> EI : Kiapp dissoc
[constants]
 Kiapp = 0.087; in μM, determine full confidence interval
[concentrations]
 E = 0.025; in μM, determine full confidence interval
[responses]
 E = 40; response per unit of enzyme, scaled to enzyme concentration (1/E)
 EI = 0
[data]
 plot logarithmic
 variable I
~~~

### Comprehensive 20059 Sequence IMPDH Phylogeny

IMPDH sequences were collected from the NCBI non-redundant protein sequence database using BLASTP with *Pbrev*IMPDHB (GenBank: ADY00133.1) as the query sequence. Non-redundant protein sequences with BLASTP E-values <10^−7^ were aligned using MAFFT (23) and a maximum likelihood phylogenetic tree was constructed using RAxML (24), using a 12-category gamma distribution for variable sites, empirically estimated amino acid frequencies, and the LG amino acid substitution matrix (25). The phylogeny (Figure S10A) was visualized with FigTree v1.4.3 software (http://tree.bio.ed.ac.uk/software/figtree/).

### Focused Eukaryotic 1371 Sequence IMPDH Phylogeny

Sequences in the eukaryotic branch of the 20059-sequence comprehensive tree were collected and duplicates were removed. The remaining 1371 sequences were aligned using MAFFT (23) and a maximum likelihood phylogenetic tree was constructed with RaxML (24). The tree (Figure S10B) was visualized with FigTree v1.4.3 software (http://tree.bio.ed.ac.uk/software/figtree/), using bacterial sequences as the outgroup.

### Fungal Phylogeny and Ancestral Sequence Reconstruction

A phylogenetic tree containing 83 fungal IMPDHs was made using the Bayesian phylogenetic software BALi-Phy (26). Outgroups were chosen from *Trichophyton*, *Microsporum*, and *Coccoidiodes* genera due to their close relation to the Eurotiale order fungi. BALi-Phy was run using three chains with the LG amino acid substitution matrix (25), site rates with discrete approximation to the Gamma distribution (27), and a symmetrical insertion-deletion model with geometrically-distributed indel lengths (28). The split posterior probabilities plateaued around 1.0 after a 20,000 iteration burn-in. The maximum *a posteriori* alignment and tree with fixed topology and branch lengths were given to IQ-Tree 2 (29), and ancestral sequences were inferred by maximum likelihood.

Ancestors to be resurrected were chosen by collating the nodes from the 80% confidence tree from the BALi-Phy run which had >80% bi-partition branch supports. Gaps in the ancestral sequences were inferred with IQ-Tree using a binary state maximum likelihood method. The alignment was recoded as 0s and 1s, corresponding to gaps and observed residues, respectively. The same maximum *a posteriori* tree topology was fixed, and branch lengths and binary ancestral gap states were inferred using the GTR binary model with binary equilibrium frequency as a free parameter. In all cases the most probable ancestral amino acid and gap state were chosen for ancestral reconstruction and resurrection. All resurrected ancestors >95% average probability across sites. Tree figures were made using the FigTree v1.4.3 software (http://tree.bio.ed.ac.uk/software/figtree/). Alignments can be downloaded at https://github.com/alexandersarkis/EvolutionOfDrugResistanceInPenicilliumFungiAWS.

### RVP inhibition assays

RVP inhibition was measured at a single concentration (1 µM) at 0.5\**K_IMP_* and 1.8\**K_NAD_*. Enzyme concentrations were set to the same concentrations as previously described, with the addition of hIMPDH2 performed at 50 nM. *K_i_* values were calculated using Equation 5:

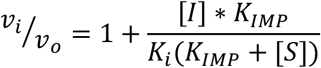

Where *v_o_* is the initial rate in the absence of inhibitor and *v_i_* is the initial rate in the presence of inhibitor. As [S] = 0.5*K_IMP_, the Equation 5 becomes:

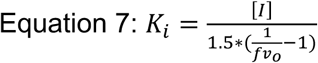

where 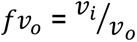.

